# Transgenerational maintenance of H3K27me3 heterochromatin is balanced by chromodomain proteins in *Caenorhabditis elegans*

**DOI:** 10.1101/2025.03.07.642071

**Authors:** Isa Özdemir, Anna Höfler, Kamila Delaney, Joanna M. Wenda, Chengyin Li, Arneet L. Saltzman, Andreas Boland, Florian A. Steiner

## Abstract

The ability to replicate and pass information to descendants is a fundamental requirement for life. In addition to the DNA-based genetic information, modifications of the DNA or DNA-associated proteins can create patterns of heritable gene regulation. Such epigenetic inheritance allows for adaptation without mutation, but its limits and regulation are incompletely understood. Here we developed a *C. elegans* system to study the transgenerational epigenetic inheritance of H3K27me3, a conserved histone posttranslational modification associated with gene repression. We find that induced alterations of the genome-wide H3K27me3 landscape and the associated fertility defects persist for many generations in genetically wildtype descendants under selective pressure. We uncover that the inheritance of the altered H3K27me3 landscape is regulated by two chromodomain proteins with antagonizing functions, and provide mechanistic insight into how this molecular memory is initiated and maintained. Our results demonstrate that epigenetic inheritance can act as a mutation-independent, heritable mechanism of adaptation.

**In Brief:** Özdemir et al. demonstrate that an altered genomic distribution of the histone modification H3K27me3 can be epigenetically inherited across many generations through the activity of HERI-1/SET-32/MES-4, which is antagonized by CEC-6/PRC2 in *C. elegans*.

**Highlights:**

- Altered H3K27me3 landscapes can be inherited for at least 15 generations in *C. elegans*.
- The chromodomain proteins CEC-6 and HERI-1 have opposite roles in antagonizing or promoting the maintenance of the altered H3K27me3 landscape.
- H3K23me3 and H3K36me3 replace H3K27me3 to promote the intergenerational and transgenerational inheritance of the altered epigenome.

## Introduction

Natural selection is based on the survival and reproduction of individuals with the best-adapted traits in a given environment, so that the genomes of those individuals are more likely to be inherited to the next generations. This process relies on the random occurrence of mutations in the genome, and is therefore slow and untargeted. However, epigenetic factors such as small RNAs, DNA methylation, or histone posttranslational modifications (PTMs) can affect gene expression in a heritable way, and can change in response to environmental influences, providing a potential mechanism for fast, targeted and reversible adaptation^1^.

Histone PTMs can act as chromatin-based epigenetic factors to influence gene expression without alterations in DNA sequence. Therefore, the extent of transmission of cellular memory by histone PTMs is one of the most-studied subjects of epigenetic inheritance (EI)^2^. There is a consensus for a semi-conservative chromatin replication model, where the parental chromatin is used as a template to replicate chromatin landscapes through the action of reader and writer proteins that recognize and catalyze histone PTMs, respectively^3^. Thus, in addition to the genetic material, the landscape of some histone PTMs is replicated in *cis* during the cell cycle. Typical examples of such “replicative EI’’ are the maintenance of repressive histone PTMs on ‘selfish’ elements^4^, or the barrier to cellular reprogramming in differentiated cell lineages^5^. However, not all histone PTMs are heritable. In cases such as the deposition of histone PTMs in response to environmental stimuli, the epigenetic remodeling in the germ line-to-soma transition, or the lineage differentiation during development, “reconstructive EI’’ takes place, where histone PTMs are targeted de novo to specific genomic regions^2,3,6–11^. The epigenome of any given cell results from both replicative EI and reconstructive EI, capacitating the germ line to give rise to highly specialized, yet totipotent germ cells^12–14^.

Examining transgenerational EI (TEI) across multiple generations is challenging due to the scarcity of a practical experimental model. For example, long generation times, mixing of epigenomes during sexual reproduction, and varying epigenetic buffering capacities among individuals pose challenges in studying TEI in mammalian systems. In contrast, self-fertilizing, isogenic *Caenorhabditis elegans* is convenient for such studies owing to the fast reproduction, invariant development and large brood sizes with little genetic variability between individuals, while at the same time sharing similar chromatin-based epigenetic systems with humans. *C. elegans* has been extensively used as a model system for the study of small RNA-based TEI, which in many cases involves changes to the histone PTMs at the targeted genomic loci^15–17^. However, whether TEI can be based directly on initial chromatin changes is less well understood^18^.

Typically, investigating EI of histone PTMs relies on the genetic removal of reader, writer or eraser proteins, and the observation of the subsequent changes in the patterns of the histone PTMs (defects in replicative EI), or the failure to initiate them de novo (defects in reconstructive EI). For instance, the polycomb histone PTM H3K27me3 is essential for germ line maintenance and cell differentiation. H3K27me3 is associated with repressed genes^19^ and can be subjected to replicative EI and reconstructive EI. Mutating polycomb group genes (PCGs) results in progressive and detrimental phenotypes such as mortal germ line (*mrt*) or maternal effect sterility (*mes*) in *C. elegans*, which are linked to the failure of maintaining or de novo initiating H3K27me3-mediated gene silencing. While such analyses allow mechanistic insight into the short-term maintenance of H3K27me3, the sterility phenotypes limit the study of H3K27me3 long-term inheritance^20,21^.

To overcome these challenges and study the long-term inheritance of H3K27me3 patterns across generations, we developed a *C. elegans* model to alter H3K27me3 landscapes by the transient expression of a mutant histone H3.3 (H3.3K27M), and then analyzed the inheritance of the altered H3K27me3 landscapes and the associated fertility defects among the wildtype offspring of the subsequent generations. *H3.3K27M* was originally identified as a dominant-negative driver mutation of human high-grade gliomas, and we previously showed that its expression in *C. elegans* leads to an autosomal loss of H3K27me3 in germ cells, resulting in aberrant gene expression, endomitotic oocytes and low fertility^22,23^. Here we showed that the H3K27me3 landscapes altered by H3.3K27M expression were epigenetically inherited for many generations of genetically wild type offspring in a metastable way. Our analysis of modifiers of this TEI identified two chromodomain proteins with antagonizing functions in the inheriting generations: CEC-6 is required to reconstruct the wildtype H3K27me3 landscape, while HERI-1 antagonizes the reconstruction of the wildtype H3K27me3 landscape by promoting the spread of ectopic H3K36me3 epialleles on autosomes. We identified an additional histone PTM, H3K23me3, that transiently replaces H3K27me3 in the first inheriting generation and mediates the initiation of H3K36me3 through the stabilization of the chromatin association of HERI-1. Together our results demonstrate that a histone PTM-based epigenetic memory can last for at least 15 generations, and we uncover a network of chromatin factors that balance the replication and reconstruction of epialleles. Our findings provide a mechanistic understanding of how organisms can transiently adapt to and inherit epigenetic changes to subsequent generations.

## Results

### Altered H3K27me3 landscapes are epigenetically inherited for many generations

We first designed an approach to transiently alter the H3K27me3 landscape in the *C. elegans* germ line, using a dominant-negative lysine 27 to methionine mutant of histone H3.3 (H3.3K27M)^22^. We expressed H3.3K27M from a tetracycline-inducible transgene present on an extrachromosomal array (*H3.3K27M^tet^*)^24^. The conditional expression of H3.3K27M and the low transmission of the extrachromosomal array to the offspring provided us with a trackable system to study the inheritance of altered H3K27me3 landscapes in genetically wildtype offspring **(Figure S1A)**.

First, we verified by immunofluorescence (IF) that *H3.3K27M^tet^*, but not wildtype worms showed a redistribution of the H3K27me3 landscape upon tetracycline treatment, with a global depletion of H3K27me3 signal on autosomes, but not on chromosome X **(Figure S1B)**. The persistence of the H3K27me3 signal on the X chromosome upon H3.3K27M^tet^ induction is explained by very low levels of H3.3 and high pre-existing levels of H3K27me3 on the X chromosome^22^. The global redistribution of H3K27me3 is easily visible by IF, which enables us to analyze the redistribution and recovery in individual cells. This provides an advantage over TEI studies using reporter genes, where the detection of chromatin changes requires ChIP-seq experiments. The *H3.3K27M^tet^*-induced changes in the H3K27me3 landscape lead to infertility at 25°C that can be classified into two sub-phenotypes: some animals are without germ line (*no gonad; resembling C. elegans PRC2 null mutants*), and the majority of them produce fewer embryos than wildtype (*subfertile; arbitrarily defined as ≤50 embryos*) **(Figure S1C)**^22^. These phenotypes become more penetrant with increasing tetracycline concentrations used for the *H3.3K27M^tet^*-induction **(Figures S1B,D)**. We selected a level of induction that caused the phenotypes, but also allowed for sufficient fertility to propagate the next generations (1 ng/µl tetracycline; **Figures S1B and S1D**). Consistent with our previous observations, *H3.3K27M^tet^*-induced infertility was temperature-sensitive **(Figure S1E)**, and was partially suppressed by the depletion of *kgb-1*, a MAPK subfamily serine-threonine kinase that is key to promoting the cell cycle activation and endomitosis observed in the germline of worms expressing H3.3K27M **(Figure S1F)**^22^.

Next, we designed a transgenerational assay to investigate the transmission of the *H3.3K27M^tet^*-induced defects to wildtype offspring. We induced the expression of *H3.3K27M^tet^*in the parental (P-1 and P0) generations. From the P0 generation, we selected offspring that had lost the *H3.3K27M^tet^* array (genetically wildtype) and no longer expressed H3.3K27M **(Figure 1A)**. In each generation, we singled 100 offspring from *fertile* individuals (selection against the inheritance of infertility; *F^fertile^*, defined as >50 viable offspring), and 50-100 offspring from *subfertile* individuals (selection for the inheritance of infertility; *F^subfertile^*, defined as ≤50 viable offspring) for further analysis. Worms that did not contain the *H3.3K27M^tet^* array in the P0 generation and therefore never expressed H3.3K27M were used as wildtype control (*P0^wt^* and *F^wt^*).

**Figure 1.**
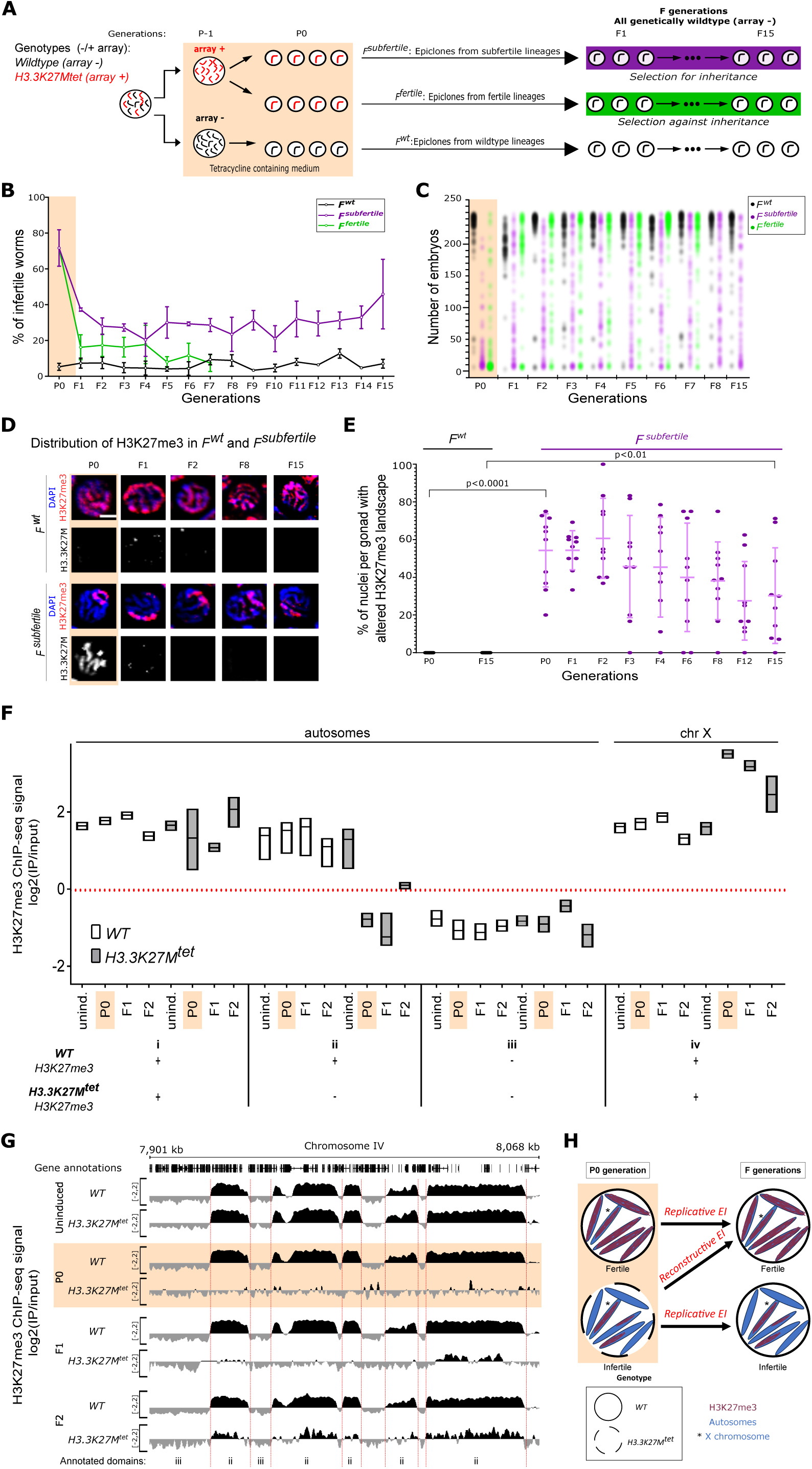
*H3.3K27M^tet^*-induced changes of the H3K27me3 landscape and the associated fertility defects are epigenetically inherited for many generations. **(A)** Experimental scheme for the study of transgenerational epigenetic inheritance of *H3.3K27M^tet^*-induced phenotypes. In the P-1 generation, wildtype and *H3.3K27M^tet^* worms were transferred onto tetracycline-containing plates. Animals carrying the *H3.3K27M^tet^*extrachromosomal array are shown in red, while black represents the loss of the array and wildtype genotype. In the P0 generation, worms were singled out on tetracycline-containing plates, and were categorized as in **Figure S1C**: *fertile* (>50 embryos) or *subfertile* (<50 embryos). Genetically wildtype worms in F generations derived from worms with *fertile* and *subfertile* phenotypes were subjected to the same categorization on plates without tetracycline. Experiments were repeated at least 3 times with 50-100 individuals per phenotype in each generation. Only P generations were exposed to tetracycline, which is represented by a beige background in all the figures. F generations are not exposed to tetracycline, and *F^subfertile^* and *F^fertile^* lineages (derived from *H3.3K27M^tet^ P0)* are highlighted with purple and green backgrounds, respectively, while *F^wt^* lineages (derived from wildtype P0) are highlighted with a white background. **(B)** Transgenerational infertility assay. Percentages of infertile worms in *F^wt^*, *F^fertile^* and *F^subfertile^* lineages from generation P0 to generation F15 at 25°C. Error bars indicate standard deviation of at least three biological replicates. **(C)** Brood size distribution of animals in *F^wt^*, *F^fertile^* and *F^subfertile^* lineages from generation P0 to generation F15 at 25°C. Each dot represents the complete brood size of an individual worm. At least 50 individuals were scored per phenotype in each generation. **(D)** Representative images from immunostaining of H3K27me3 and OLLAS-tagged H3.3K27M in late pachytene cells in generations P0, F1, F2, F8 and F15 of *F^wt^* and *F^subfertile^*. Scale bar represents 2 µm. **(E)** Percentages of pachytene nuclei with altered H3K27me3 landscape per gonad in *P0^wt^ and F15^wt^* and *P0-F15^subfertile^*. Each dot represents the percentage of pachytene nuclei with altered H3K27me3 patterns within an individual gonad (at least 10 nuclei per gonad). At least 10 gonads were scored per generation, and means with standard deviations are shown. P-values are from 2-way ANOVA comparisons. n.s. stands for not significant. *F2-14^wt^* generations were identical to *P0^wt^* and *F15^wt^ (*not shown). **(F)** Average H3K27me3 ChIP-seq signal genome-wide for each domain type from uninduced (unind.), tetracycline-induced P0, and F1 and F2 wildtype (WT) and *H3.3K27M^tet^* strains at 25°C. Domains were annotated based on H3K27me3 occupancy changes between wildtype and *H3.3K27M^tet^* in the P0 generation as in^22^. i autosomal H3K27me3 signal is preserved, ii autosomal H3K27me3 signal is depleted (this includes the vast majority of the autosomal H3K27me3 regions), iii autosomal H3K27me3 signal is not detected in *F^wt^* and *F^subfertile^*, and iv is chromosome X. **(G)** Genome browser views of H3K27me3 ChIP-seq experiments from uninduced, tetracycline-induced P0, and F1 and F2 wildtype (WT) and *H3.3K27M^tet^* strains at 25°C. A representative part of chromosome IV (∼160 kb) is shown. Each ChIP-seq track is the average log_2_ ratio (IP/input) of three independent biological replicates. The borders between annotated domains are shown with red dashed lines. Domain types were defined as in Figure 1F. **(H)** Cartoon of pachytene nuclei showing EI of *H3.3K27M^tet^-*induced changes in H3K27me3 distribution from P0 to the inheriting F generations. The *H3.3K27M^tet^*expressing pachytene nucleus is shown with a dashed border, and genetically wildtype nuclei are shown with solid borders. Chromosomes are shown in blue, and the X chromosome is marked by an asterisk. The H3K27me3 distribution is shown in magenta. Expression of *H3.3K27M^tet^* initiates the chromatin changes and the associated fertility defects. During the transition from one generation to the next, some germ cells reconstruct the wildtype epigenetic state (reconstructive EI), leading to fertility, while other germ cells replicate the parental epialleles (replicative EI), leading to inheritance of the infertility phenotypes.

In the P0 generation, 70% of the worms carrying the *H3.3K27M^tet^* array showed fertility defects, while less than 8% of the *P0^wt^* worms were subfertile at 25°C **(Figure 1B)**. Remarkably, the *H3.3K27M^tet^*-induced fertility defects were still present among the genetically wildtype F1 offspring. About 40% of the *F1^subfertile^* and about 20% of the *F1^fertile^*inherited their parental fertility defects **(Figure 1B)**. We expanded the TEI assays to generations F2 to F15. We found that when selecting for *subfertile* lineages in each generation, the fertility defects were present in about 30-40% of the worms of the following generation. This demonstrates that under selection, the consequences of alterations in H3K27me3 landscape in the P0 generation can be heritable for at least 15 generations. The fertility defects among the *F^fertile^* lineages were less pronounced and became comparable to the *F^wt^* at generation F5 **(Figure 1B)**. The offspring of *subfertile* worms displayed each sub-phenotype (*no gonad*, *subfertile* or *fertile*) in each generation, implying that the *H3.3K27M^tet^*-induced changes in the H3K27me3 landscape are either replicated, or the wildtype H3K27me3 landscape is reconstructed, resulting in variability in the fertility phenotypes of the next generation **(Table S1)**. To obtain a quantitative measure of the fertility defects, we counted the number of embryos laid by the *F^wt^*, *F^fertile^* and *F^subfertile^* worms **(Figure 1C; Table S2)**. The offspring derived from *F^wt^* animals laid >200 embryos in each generation, whereas the brood size of *H3.3K27M^tet^*-induced *P0* animals was strongly reduced. The *F^fertile^* descendants had brood sizes close to wildtype after one generation, with slightly increased variability. The brood sizes of *F^subfertile^* were on average consistently lower for at least 15 generations, with a large variability in every generation, indicating that some worms recovered wild-type brood sizes, while others inherited the fertility defects **(Figure 1C; Table S2)**.

Next, we tested whether the inheritance of fertility defects was based on the inheritance of the altered H3K27me3 landscape. We used IF to detect H3K27me3 in germ cells of *F^wt^* and *F^subfertile^*worms in generations P0 to F15. IF allows the detection of altered chromatin landscapes within single cells and is therefore ideally suited to probe patterns that are inherited with low penetrance and are heterogeneous within individuals. *H3.3K27M^tet^ P0* worms showed germ cells with an altered H3K27me3 landscape, where the H3K27me3 signal is depleted from autosomes but retained on chromosome X. The almost complete autosomal loss of H3K27me3 facilitated the identification of cells with aberrant H3K27me3 landscapes in the offspring and allowed the quantification of the penetrance of the inheritance. Strikingly, the changed H3K27me3 landscape was present in the next 15 *F* generations, even though the H3.3K27M protein was no longer detectable in any of the *F* generations **(Figures 1D and S2A)**. These results demonstrate that both the altered H3K27me3 landscape and the resulting fertility defects induced by *H3.3K27M^tet^*in the *P0* generation can be inherited for at least 15 generations in a genetically wildtype population. To assess the penetrance of the inherited chromatin changes in *F^subfertile^* animals, we counted the percentage of nuclei with clearly recognizable altered H3K27me3 patterns per gonad **(Figure 1E)**. Most *F^subfertile^* individuals contained a mix of germ cells with replicated altered H3K27me3 landscapes and fully or partially reconstructed wildtype-like H3K27me3 landscapes. We only counted nuclei with fully replicated defects as “altered” **(Figure S2B)**. *F^wt^* individuals did not show any altered patterns and faithfully replicated the wildtype H3K27me3 landscapes. There was considerable variability between *F^subfertile^* individuals **(Figure S2B)**, and the frequency of nuclei per gonad with altered H3K27me3 landscapes decreased with increasing generation number **(Figure 1E)**. Despite this variability, some individuals still had up to 70% germ cells with altered H3K27me3 landscapes in generation F15, demonstrating that they can be robustly transmitted across generations **(Figure 1E)**. The variability among germ cells explains how individuals in each generation can give rise to both *fertile* and *subfertile* offspring.

To verify that H3K27me3 was reconstructed within the same genomic domains where it was lost upon *H3.3K27M^tet^* induction, we assessed the H3K27me3 landscapes by ChIP-seq in P0, F1 and F2 populations, with uninduced and wildtype strains as controls **(Figures 1F,G and S2C)**. The ChIP-seq experiments are not quantitative, but allow a relative comparison between the autosomal and X-chromosomal levels. For the analysis, we divided the genome into four types of domains, as done in our previous publication^22^: i autosomal H3K27me3 signal is preserved upon *H3.3K27M^tet^* induction (these only cover a small fraction of the genome), ii autosomal H3K27me3 signal is depleted upon *H3.3K27M^tet^* induction (these include the vast majority of autosomal H3K27me3 domains), iii autosomal H3K27me3 signal is not detected in any condition, iv chromosome X. Consistent with the observations by IF, the majority of autosomal **(Figures 1F,G; domains ii)**, but not X-chromosomal H3K27me3 domains **(Figures 1F,G; domains iv)**, were erased in the *H3.3K27M^tet^*-induced P0 animals compared to wildtype and uninduced controls. We observed a gradual reconstruction of the wildtype H3K27me3 landscape in the F generations, which was restricted to the domains with H3K27me3 signal in the wildtype **(Figures 1F,G; domains ii).** These observations are consistent with a reconstruction of the original H3K27me3 landscape in part of the germ cells, which are averaged with the replicated altered H3K27me3 landscapes in other germ cells in the F1 and F2 generations. Overall, we observed a reconstruction of the wildtype H3K27me3 landscape in some of the offspring, yet a considerable portion of offspring replicated the *H3.3K27M^tet^*-induced changes **(Figure 1H).**

### CEC-6 is required for the reconstruction of the wildtype H3K27me3 landscape in the F generations

Next, we aimed to identify factors that regulate the reconstruction of the wildtype and the replication of the altered H3K27me3 landscapes during TEI in *F^subfertile^* lineages. We reasoned that depletion of factors required for replicative TEI would result in a loss of the heritable phenotypes, and the depletion of factors involved in reconstructive TEI would increase the penetrance of the heritable phenotypes. We focused on *C. elegans* chromodomain (CEC) proteins^25^, as they were previously shown to play a role in TEI^26,27^. CEC proteins are homologs of CBX and Pc proteins, which are readers of methylated lysine residues on histone H3 in mammals and *Drosophila*^28,29^.

We first examined the role of CEC-6, which has been identified as a reader of H3K27me2/3 in the *C. elegans* germ line^26,30^. *cec-6 KO* animals show a *mortal germ line (mrt)* phenotype at 22°C after ∼40 generations, suggesting that it has a role in the maintenance of H3K27me3 across generations^26^. We analyzed the TEI of *H3.3K27M^tet^*-induced defects either in *cec-6 RNAi* or *cec-6 KO* background **(Figures 2A, S3A,B)**. Elimination of CEC-6 did not affect the fertility of *F^wt^*animals in the generations analyzed. It also did not modify the fertility defects in the *H3.3K27M^tet^* P0 generation. However, it enhanced the inheritance of fertility defects in *F^subfertile^* lineages, and we now observed around 70% infertile animals (compared to 30-40% in wildtype background) in the *F1^subfertile^* and *F2^subfertile^*generations **(Figures 2A and S3B)**. The *H3.3K27M^tet^*-induced changes of the H3K27me3 landscapes in *P0* and *F^subfertile^* animals were not affected by the loss of CEC-6, but the frequency of germ cells with an altered H3K27me3 landscape increased, especially in the *F1^subfertile^* and *F2^subfertile^*generations, consistent with the enhanced penetrance of the fertility defects **(Figures 2A-C)**. The persistence of the H3K27me3 signal on the X chromosome in *cec-6* KO worms suggests that CEC-6 is not required for the maintenance of H3K27me3 domains per se. Instead, the increased frequency of germ cells with altered H3K27me3 patterns suggests that CEC-6 is required for the reconstruction of wildtype H3K27me3 domains that have been lost due to *H3.3K27M^tet^* expression in the P0 generation. *cec-6 KO* alone also resulted in a gradual loss of wildtype H3K27me3 epialleles in some *F2^wt^* worms, which is consistent with the previously observed progressive *mrt* phenotype of *cec-6* mutants **(Figure 2C)**^26^. In contrast, depletion of the somatic H3K27me2/3 reader *cec-1*^26^ did not result in any fertility defects **(Figure S3B)**, suggesting that the TEI of the fertility defects is germline-based, and not caused by somatic defects in the inheriting generations.

**Figure 2.**
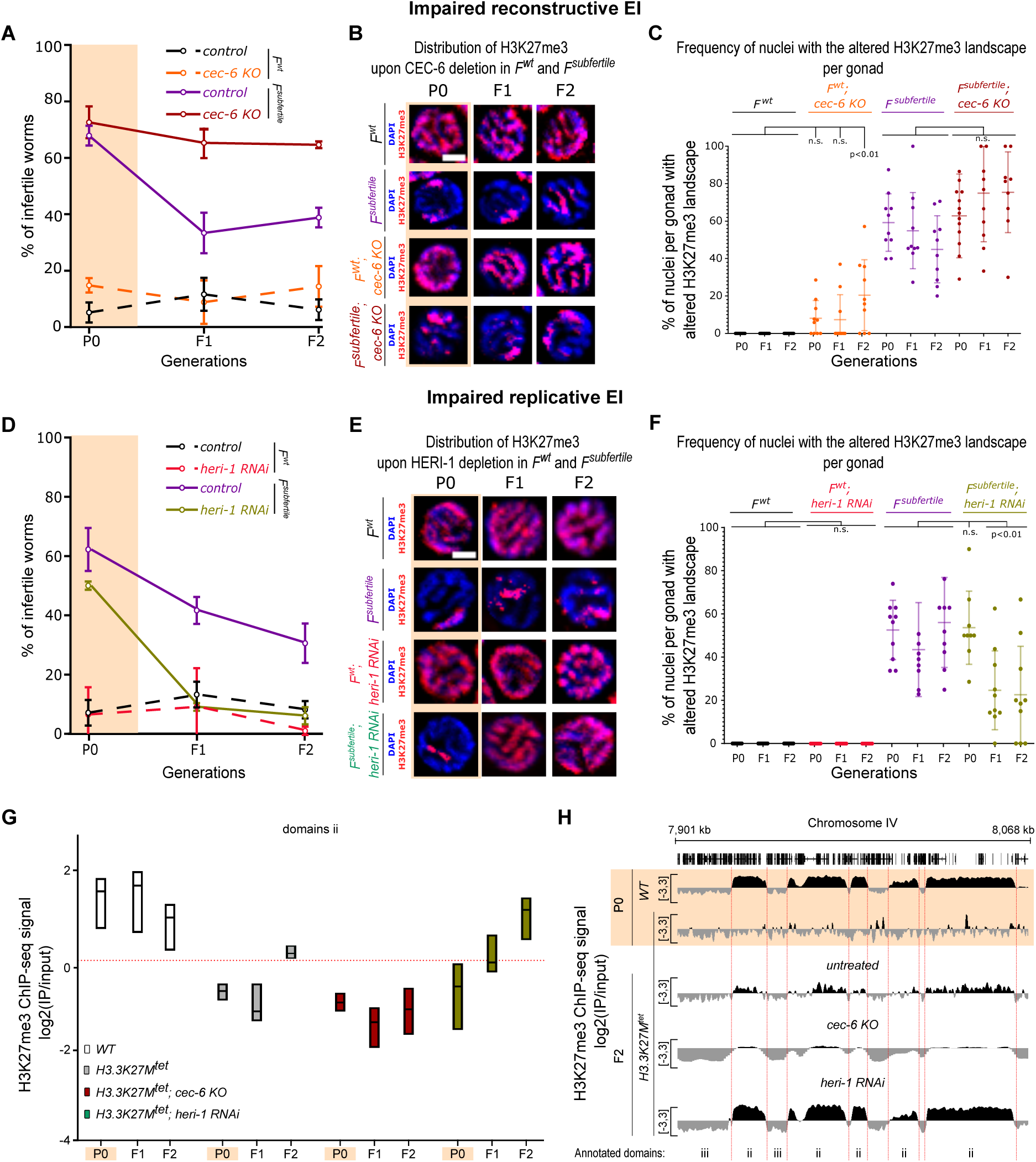
CEC-6 is essential for the reconstruction of the wildtype H3K27me3 domains while HERI-1 is required for replicative EI of the *H3.3K27M^tet^*-induced defects. *P0-F2^wt^* and *P0-F2^subfertile^* generations grown at 25°C were analyzed in each panel, as indicated. **(A-C)** Comparison of wildtype and *cec-6 KO*. **(A)** Transgenerational infertility assay. Percentages of infertile worms in *P0-F2^wt^* (dashed lines) and *P0-F2^subfertile^* (solid lines) generations. **(B)** Representative images from immunostaining of H3K27me3 in late pachytene nuclei. **(C)** Frequency of pachytene nuclei with altered H3K27me3 landscape. **(D-F)** Comparison of *control RNAi* and *heri-1 RNAi*. **(D)** Transgenerational infertility assay. Percentages of infertile worms in *P0-F2^wt^* (dashed lines) and *P0-F2^subfertile^* (solid lines) generations. **(E)** Representative images from immunostaining of H3K27me3 in late pachytene nuclei. **(F)** Frequency of pachytene nuclei with altered H3K27me3 landscape. Error bars indicate standard deviations of at least three biological repeats in **(A, D)**. Scale bars represent 2 µm in **(B, E)**. In **(C, F)** each dot represents the percentage of pachytene nuclei with altered H3K27me3 patterns within an individual gonad (at least 10 nuclei per gonad). At least 10 gonads were scored per generation, and means with standard deviations are shown. P-values are from 2-way ANOVA with Sidak’s multiple comparison tests. n.s. stands for not significant. **(G)** Average H3K27me3 ChIP-seq signal genome-wide for type ii domains (autosomal H3K27me3 domains in wildtype; as in Figure 1F) in wildtype (WT) and *H3.3K27M^tet^* with or without CEC-6 or HERI-1. The WT P0-F2 (white) and *H3.3K27M^tet^*P0-F2 (gray) bars are reproduced from Figure 1F. **(H)** Genome browser views of H3K27me3 levels as determined by H3K27me3 ChIP-seq experiments in the presence of both CEC-6 and HERI-1 (untreated), or in *cec-6 KO* background or upon *heri-1 RNAi*. The top three tracks are reproduced from Figure 1G. A representative part of chromosome IV (∼160 kb) is shown (same as in Figure 1G). Each ChIP-seq track is the average log_2_ ratio (IP/input) of three independent biological replicates. The borders between annotated domains are shown with red dashed lines. Domain types were defined as in Figure 1F.

### HERI-1 is required for the replication of altered H3K27me3 epialleles in the F generations

We next investigated the role of HERI-1 in the inheritance of H3K27me3 landscape. HERI-1 is a CEC protein that was identified in a genetic screen for factors limiting RNAi-induced TEI^27,30^. We analyzed the TEI of *H3.3K27M^tet^*-induced defects upon *heri-1 RNAi* **(Figures 2D and S3A)**. *heri-1* RNAi did not affect the fertility of P0 and *F^wt^* worms. However, contrary to *cec-6 KO*, *heri-1 RNAi* fully suppressed the inheritance of fertility defects in *F1^subfertile^***(Figure 2D)**. Detection of H3K27me3 by IF revealed that *heri-1 RNAi* promoted the reconstruction of H3K27me3 epialleles in most of the *F^subfertile^* lineages, while it neither changed the H3K27me3 landscape in *F^wt^*, nor prevented the chromatin defects in *H3.3K27M^tet^* P0 generation **(Figures 2E,F)**. These observations suggested that HERI-1 is involved in the replicative EI of the *H3.3K27M^tet^*-induced defects and counteracts the reconstruction of the wildtype H3K27me3 landscape by CEC-6 in *F^subfertile^* lineages **(Figure S3C)**.

Analysis of H3K27me3 by ChIP-seq confirmed that *cec-6 KO* or *heri-1 RNAi* promoted replicative EI or reconstructive EI of wildtype H3K27me3 epialleles, respectively, in the F generations **(Figures 2G,H)**, and thus support the interpretation of the IF analysis. To explore the antagonism between CEC-6-mediated reconstructive EI and HERI-1-mediated replicative EI, we simultaneously removed CEC-6 and HERI-1 in the TEI assay **(Figure S3D)**. The double depletion resulted in rebalancing of the inheritance of the fertility defects that phenocopied *F^subfertile^* in wildtype background, suggesting that CEC-6-mediated reconstructive EI and HERI-1-mediated replicative EI function independently of each other.

### Chromatin association of CEC-6 and HERI-1 is required for their roles in TEI

Given strong sequence similarities of the chromodomains of CEC-6 and HERI-1 and the similar N-terminal positions of the chromodomains within the proteins **(Figures S4A,B),** the opposing effects on the TEI of the H3K27me3 landscape in *F^subfertile^* lineages were intriguing. The chromodomain structures of CEC-6 and HERI-1 predicted by Alphafold2^31,32^ both possess three twisted antiparallel beta-strands followed by an alpha-helix, which resembles the chromodomain of *Polycomb* (Pc), the reader for H3K27me2/3 in *Drosophila* **(Figure S4C)**^28,29^. To investigate the antagonistic roles of CEC-6 and HERI-1, we followed their localization during TEI by IF. In *F^wt^* animals, CEC-6 was mainly chromatin-bound, and showed 90% colocalization with H3K27me3 **(Figure 3A)**. Upon *H3.3K27M^tet^*-induced changes in H3K27me3 landscape in the P0 generation, this colocalization was reduced to about 40%, and remained low in *F^subfertile^* **(Figure 3A).** In germ cells of *F^subfertile^*animals, CEC-6 occupied chromosomal regions without H3K27me3, supporting the model that CEC-6 functions in pioneering the reconstruction of H3K27me3 domains. HERI-1 showed a different nuclear distribution compared to CEC-6. In *F^wt^* animals, HERI-1 was diffusely distributed in the nucleoplasm. Upon *H3.3K27M^tet^*-induced changes in the H3K27me3 landscape, it switched from this diffuse pattern to a chromatin-associated pattern, but showed low colocalization with H3K27me3 and remained excluded from the X chromosome, suggesting that it was only targeted to regions where H3K27me3 was depleted **(Figure 3B)**. This pattern was maintained in *F^subfertile^* lineages **(Figure 3B)**. These observations are consistent with the observed role of HERI-1 in antagonizing the reconstruction of the wildtype H3K27me3 landscape and thus in replicating the *H3.3K27M^tet^*-induced changes of the H3K27me3 landscape **(Figure 3C)**. The localization of CEC-6 and HERI-1 to regions of the nuclei lacking H3K27me3, together with the observed phenotypes upon depletion of these chromodomain proteins, suggest a model where both CEC-6 and HERI-1 target the autosomal regions that become depleted from H3K27me3, where they have opposite roles: CEC-6 acts in reconstructing the H3K27me3 domains, while HERI-1 antagonizes this reconstruction.

**Figure 3.**
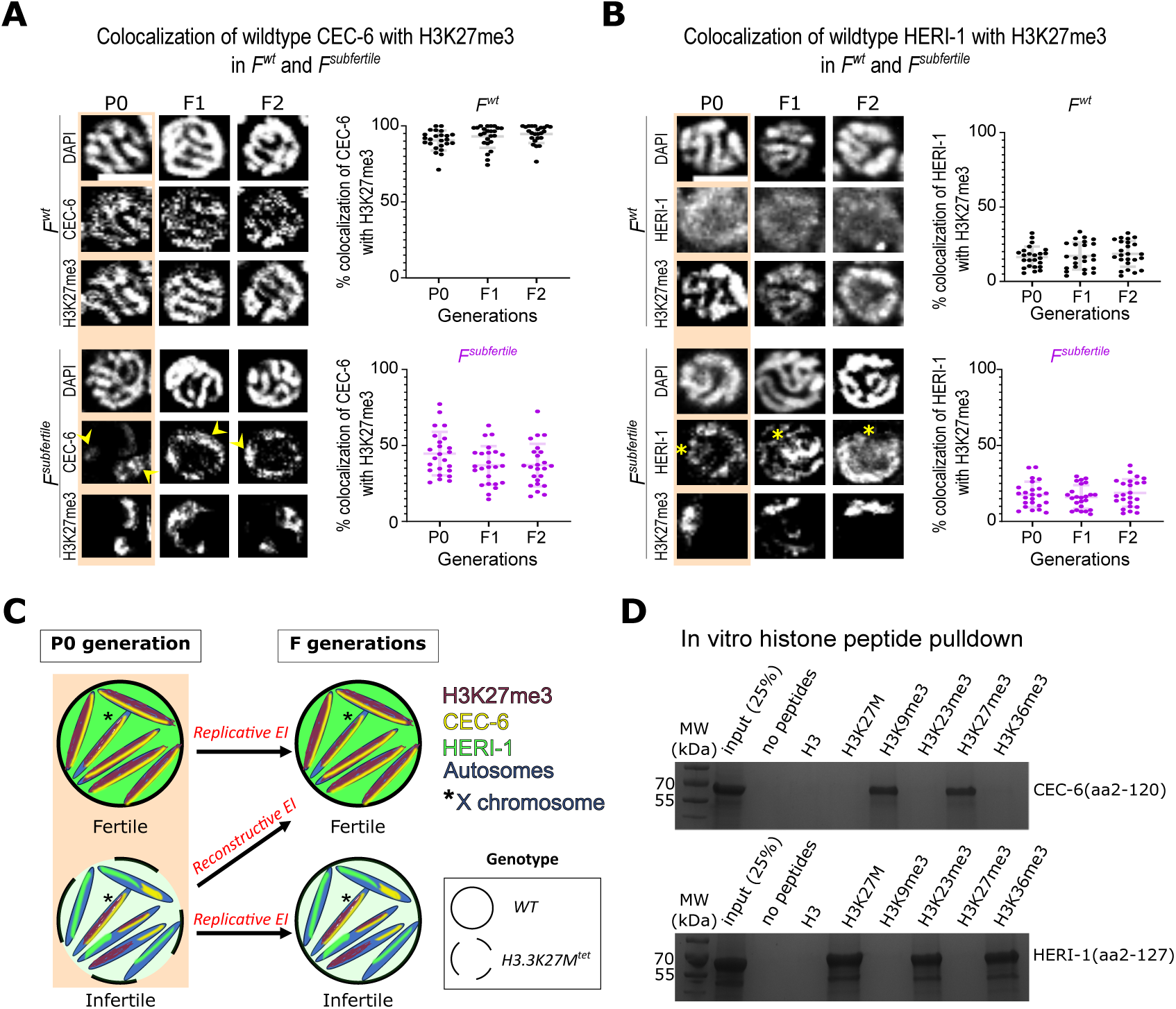
CEC-6 and HERI-1 show distinct chromatin dispositions. **(A, B)** Representative images from immunostaining of H3K27me3 and FLAG-tagged CEC-6 or HERI-1 in late pachytene nuclei of *P0-F2^wt^* and *P0-F2^subfertile^* generations, and quantification of colocalization of chromodomain proteins with H3K27me3. At least 20 nuclei were scored per generation, and means with standard deviation are shown. Scale bars represent 4 µm. Arrowheads point to CEC-6 bound to H3K27me3-free chromatin in **(A)**, asterisks indicate positions of H3K27me3-enriched X chromosomes in **(B)**. **(C)** Cartoon summarizing the sub-nuclear localization of CEC-6 (yellow) and HERI-1 (green) in pachytene nuclei of *F^wt^* and *F^subfertile^*. The *H3.3K27M^tet^*expressing pachytene nucleus is shown with a dashed border, and genetically wildtype nuclei are shown with solid borders. Chromosomes are shown in blue, and the X chromosome is marked by an asterisk. The H3K27me3 distribution, CEC-6 and HERI-1 are shown in magenta, yellow and green, respectively. Expression of *H3.3K27M^tet^* induced heritable chromatin localization changes of CE6 and HERI-1. CEC-6 was chromatin bound in both *F^wt^* and *F^subfertile^* conditions while HERI-1 was nucleoplasmic in *F^wt^* and mostly chromatin-bound in *F^subfertile^*. **(D)** A Coomassie staining of CEC-6(aa2-120) and HERI-1(aa2-127) after in vitro histone peptide pull-down.

Chromodomains of Pc-like proteins form an aromatic cage to bind to methylated lysine residues^28,29^. We next tested if this methyl-lysine binding activity is important for the roles of CEC-6 and HERI-1 in regulating TEI of H3K27me3 domains. We mutated two of the conserved cage-forming aromatic residues in the linker region of the beta2 and beta3 strands (*cec-6(W55A, Y58A)* and *heri-1(W62A, Y65A)*; **Figures S4B,C**), which should abolish the binding to methylated lysines. Introduction of these point mutations phenocopied the *cec-6 KO* and *heri-1 RNAi*, respectively: they did not affect the (in)-fertility of *H3.3K27M^tet^*P0 or *F^wt^*, but abolished the reconstructive EI and replicative EI, respectively, in the *F^subfertile^* generations **(Figures S5A,B)**. Detection of CEC-6(W55A, Y58A) and HERI-1(W62A, Y65A) by IF revealed the loss of chromatin binding of these chromodomain mutant proteins in both *F^wt^* and *F^subfertile^*animals **(Figures S5C,D)**. These results demonstrate that the chromodomains of CEC-6 and HERI-1 are required for their chromatin localization, and that the binding of these proteins to chromatin is required for their role in reconstructing or replicating chromatin states.

### CEC-6 binds to H3K27me3 and H3K9me3 whereas HERI-1 binds to H3K27M, H3K36me3 and H3K23me3

Both CEC-6 and HERI-1 change their chromatin localization upon expression of *H3.3K27M^tet^*. We therefore wondered if these chromodomain proteins showed any binding specificity towards the H3.3K27M mutant or specific histone PTMs. We purified recombinant versions of the chromodomains of CEC-6 (aa2-120) and HERI-1 (aa2-127), mixed them with biotinylated, modified or unmodified histone H3 peptides, and tested for co-precipitation in vitro **(Figures S6A-C)**. The CEC-6 chromodomain has previously been shown to bind to di-/trimethylated H3K9 and H3K27 in a histone peptide array screen^26^. The binding of the HERI-1 chromodomain has not been tested in vitro, but HERI-1 was shown to be recruited to RNAi-targeted genomic loci, which required SET-32, a germline-specific HMT for H3K23^15,27,33,34^. In our in vitro histone peptide pulldown assay, we therefore included unmodified H3, H3K27M, H3K9me3, H3K27me3 and H3K23me3, as well as H3K36me3, an antagonistic mark to H3K27me3^35^. We found that CEC-6(2-120) specifically recognized the H3K9me3 and H3K27me3 peptides, consistent with previous findings^26,30^, while HERI-1(2-127) showed affinity to the H3K27M, H3K23me3, and H3K36me3 peptides **(Figures 3D, S6D,E)**. These results suggest that CEC-6 binds to heterochromatin and likely acts in concert with PRC2 to reconstruct wildtype H3K27me3 domains, whereas HERI-1 is initially recruited to chromatin by H3K27M in P0, and promotes replicative EI through binding to H3K36me3 and/or H3K23me3 in *F^subfertile^* generations.

### Spreading of H3K36me3 promotes TEI of altered H3K27me3 patterns

Given the affinity of HERI-1 to H3K36me3, we tested whether H3K36me3 was required for the replicative EI of *H3.3K27M^tet^*-induced changes in the H3K27me3 landscape. H3K27me3 and H3K36me3 form mutually exclusive domains across the genome in the *C. elegans* germ line^20^. We therefore hypothesized that the H3K27me3 domains that are lost upon expression of H3K27M are replaced by ectopic H3K36me3 deposition. These H3K36me3 epialleles might then antagonize the reconstruction of wildtype H3K27me3 epialleles during TEI. Co-staining of H3K36me3 and H3K27me3 in worms from *P0* to *F8^subfertile^* revealed an autosomal spread of H3K36me3 in pachytene nuclei where H3K27me3 was reduced **(Figure 4A)**. However, the H3K36me3 increase was not significant in the *P0* generation, and only became penetrant in generations after *F1^subfertile^* **(Figures 4A,B).** The spread of H3K36me3 patterns revealed a considerable variability in each subsequent generation due to varying competition between replicative EI and reconstructive EI **(Figure S7)**.

**Figure 4.**
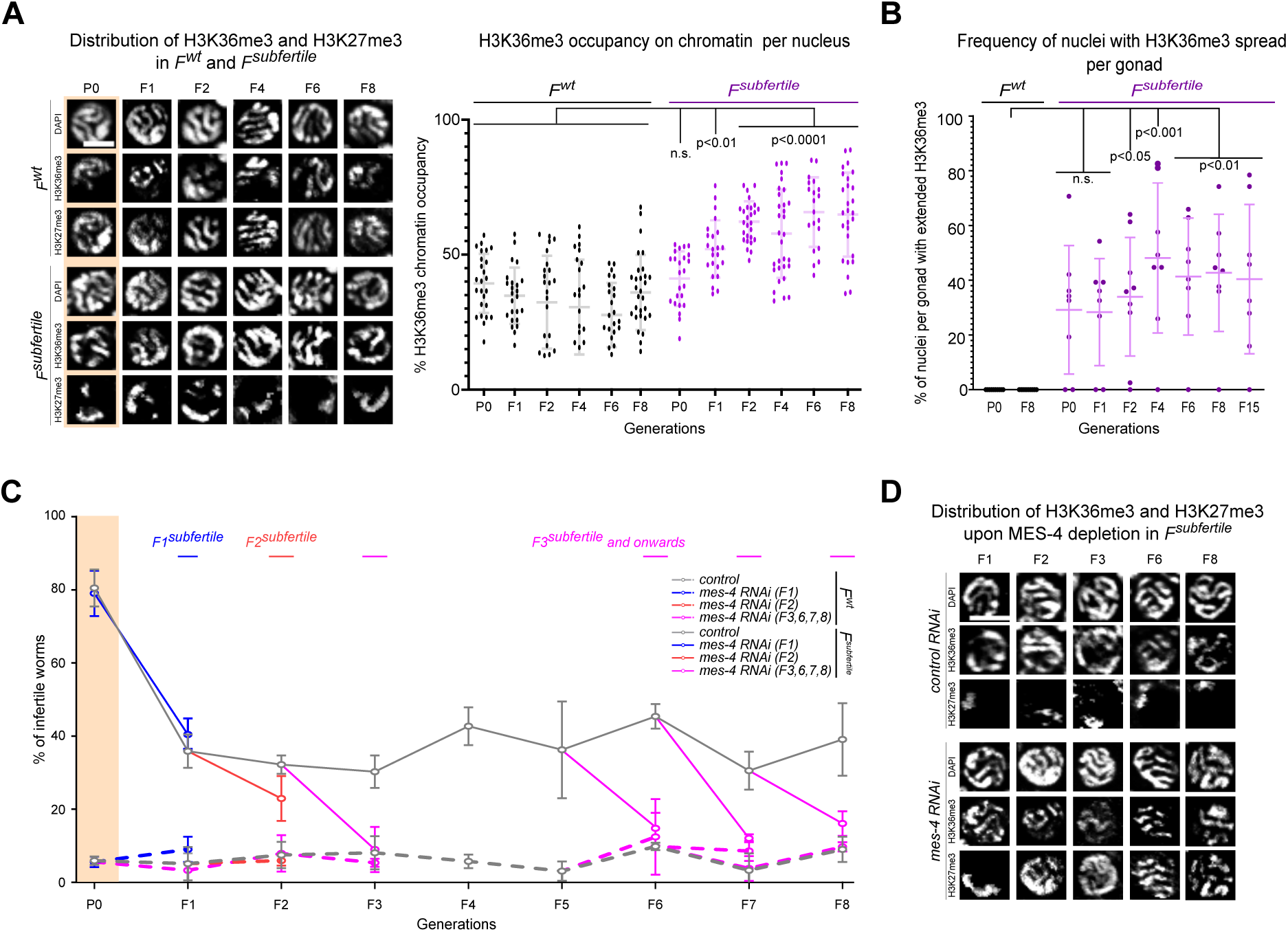
The spread of H3K36me3 is required for replicative EI of *H3.3K27M^tet^-induced* phenotypes in generations after *F2^subfertile^*. **(A)** Representative images from immunostaining of H3K27me3 and H3K36me3 in late pachytene nuclei, and quantification of H3K36me3 chromatin occupancy, in *P0-F8^wt^* and *P0-F8^subfertile^* generations. In the quantification panel, each dot represents the percent of chromatin with H3K36me3 signal per nucleus. At least 20 nuclei were scored per generation, and means with standard deviations are shown. **(B)** Frequency of pachytene nuclei with extended H3K36me3 signal in *P0/F8^wt^* and *P0-F8^subfertile^* generations. Each dot represents the percentage of pachytene nuclei with extended H3K36me3 signal within an individual gonad (at least 10 nuclei per gonad). At least 7 gonads were scored per generation, and means with standard deviations are shown. *F2-F7^wt^*generations were identical to *P0/F8^wt^ (*not shown). P-values in **(A and B)** are from 2-way ANOVA with Sidak’s multiple comparison tests in comparison to corresponding *F^wt^* and *F^subfertile^*generations. n.s. stands for not significant. **(C)** Transgenerational infertility assay. Percentages of infertile worms in *P0-F8^wt^* (dashed lines) and *P0-F8^subfertile^* (solid lines) generations, with *control RNAi* (gray) or *mes-4 RNAi* treatment in F1 (blue), F2 (red), F3, F6, F7 or F8 (fuchsia) at 25°C. Means with standard deviations of at least three biological replicates are shown. **(D)** Representative images from immunostaining of H3K27me3 and H3K36me3 in late pachytene nuclei of *F1-F8^subfertile^*generations with *control RNAi* or *mes-4 RNAi* treatment. Scale bars represent 4 µm in **(A and D)**.

To test if the spreading of H3K36me3 is required for the replicative TEI of the *H3.3K27M^tet^*-induced defects, we depleted MES-4, the histone methyltransferase required for the maintenance of H3K36me3^20,36,37^. Loss of MES-4 results in a maternal-effect sterile *(mes)* phenotype^37^, and we could therefore only analyze the generation immediately following *mes-4 RNAi*. *mes-4 RNAi* in *F2^subfertile^*or later generations suppressed the replicative EI of *F^subfertile^* defects, and worms of the following generation showed fertility and H3K27me3 landscapes comparable to wildtype **(Figures 4C,D)**. The suppression of the fertility defects upon *mes-4 RNAi* demonstrates that the ectopic H3K36me3 epialleles formed upon the expression of *H3.3K27M^tet^* prevent the reconstruction of H3K27me3 domains and promote the replicative TEI of the *H3.3K27M^tet^*-induced defects. However, *mes-4* depletion had no suppressive effect on the inherited fertility defects or on the altered H3K27me3 landscape in *F1^subfertile^* worms, and an only intermediate suppressive effect on the *F2^subfertile^* worms **(Figures 4C,D)**. These results suggest that MES-4 is required for maintaining the ectopically initiated H3K36me3 epialleles in *F^subfertile^*, but not for initiating the TEI during the transition from P0 to *F1-F2^subfertile^*. **H3K23me3 is required for the initiation of TEI.** The depletion of MES-4 suppressed the replicative TEI in generations after *F2^subfertile^*, while the depletion of HERI-1 suppressed the replicative TEI of *H3.3K27M^tet^*-induced defects from P0 to *F2^subfertile^* **(Figures 2D-F)**. Given the physical interaction between HERI-1 and H3K23me3 **(Figure 3D),** and the SET-32-dependent localization of HERI-1 to chromatin^27^, we hypothesized that H3K23me3 initially replaces H3K27me3 and is required for initiating the TEI of the *H3K27M^tet^*-induced defects. To test this hypothesis, we detected H3K23me3 by IF from P0 to F8 in *F^wt^* and *F^subfertile^* animals. Consistent with previous observations^38^, H3K23me3 appeared as localized spots on chromatin in *F^wt^* animals **(Figure 5A)**. Upon the induction of *H3.3K27M^tet^*, it appeared as a continuous pattern on autosomes in P0 and *F1-F2^subfertile^* generations **(Figure 5A)**. However, the frequency of pachytene nuclei with this increased H3K23me3 occupancy decreased with each generation, and mainly the wildtype distribution was observed after *F2^subfertile^* **(Figure 5B)**.

**Figure 5.**
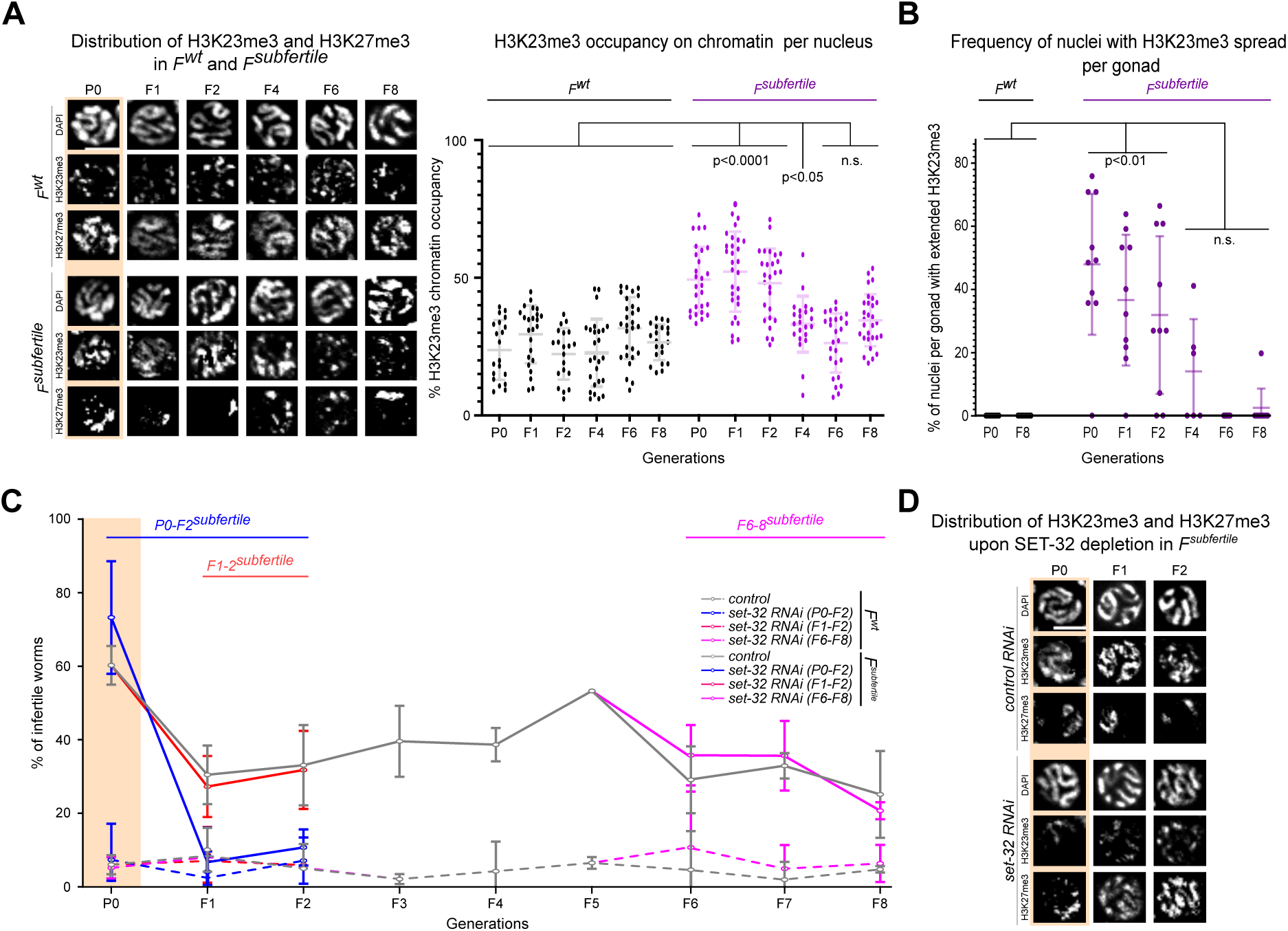
H3K23me3 replaces H3K27me3 in early generations to initiate replicative EI of ***H3.3K27M^tet^-*induced defects**. **(A)** Representative images from immunostaining of H3K27me3 and H3K23me3 in late pachytene nuclei, and quantification of H3K23me3 chromatin occupancy, in *P0-F8^wt^* and *P0-F8^subfertile^* generations. In the quantification panel, each dot represents the percent of chromatin with H3K23me3 signal per nucleus. At least 20 nuclei were scored per generation, and means with standard deviations are shown. **(B)** Frequency of pachytene nuclei with extended H3K23me3 signal in *P0/F8^wt^* and *P0-F8^subfertile^* generations. Each dot represents the percentage of pachytene nuclei with extended H3K23me3 signal within an individual gonad (at least 10 nuclei per gonad). At least 10 gonads were scored per generation, and means with standard deviations are shown. *F2-F7^wt^*generations were identical to *P0/F8^wt^ (*not shown). P-values in **(A, B)** are from 2-way ANOVA with Sidak’s multiple comparison tests in comparison to corresponding *F^wt^* and *F^subfertile^*generations. n.s. stands for not significant. **(C)** Transgenerational infertility assay. Percentages of infertile worms in *P0-F8^wt^* (dashed lines) and *P0-F8^subfertile^* (solid lines) generations, with *control RNAi* (gray) or *set-32 RNAi* treatment from P0 to F2 (blue) or from F1 to F2 (red) or from F6 to F8 (fuchsia) at 25°C. Means and standard deviations of at least three biological replicates are shown. **(D)** Representative images from immunostaining of H3K27me3 and H3K23me3 in late pachytene nuclei of *P0-F2^subfertile^*generations with *control RNAi* or *set-32 RNAi* treatment. Scale bars represent 4 µm in **(A and D)**.

We next examined whether the increased deposition of H3K23me3 was required for the replicative TEI of the *H3.3K27M^tet^*-induced phenotypes, and whether it could bridge the reduction of H3K27me3 and the increase of H3K36me3 during the P0 to *F1^subfertile^* transition. SET-32, the main HMT for H3K23me3 in the germ line **(Figure S8A)**, showed a diffuse pattern in wildtype and was targeted to chromatin in *H3.3K27M^tet^* P0 **(Figure S8B)**, similar to the dynamics observed for HERI-1 **(Figure 3B)**. However, unlike HERI-1, the chromatin association of SET-32 was lost again in *F1^subfertile^*, where SET-32 resumed a diffuse pattern as in wildtype **(Figure S8B)**. This implies that the expression of *H3.3K27M^tet^*induces the chromatin localization of SET-32, leading to the deposition of H3K23me3 in the P0 generation, and that those ectopic H3K23me3 epialleles counteract the reconstructive EI of wildtype H3K27me3 domains in the following generation. To test this hypothesis, we examined the inheritance of *H3.3K27M^tet^*-induced fertility defects in worms without SET-32 activity, either by *set-32 RNAi* or by using a s*et-32(red11)* loss-of-function allele^16^. Loss of SET-32 activity phenocopied the loss of HERI-1 in early generations: the *H3K27M^tet^-*induced fertility defects were unchanged in the P0, but fully suppressed in the F1 and F2 generations and restored the wildtype H3K27me3 landscape in *F1-F2^subfertile^* **(Figures 5C,D and S8C)**. However, *set-32 RNAi* in *F1-F2^subfertile^* or *F6-8^subfertile^* did not suppress the inheritance of fertility defects **(Figure 5C)**. These results suggested that SET-32 plays a major role in the intergenerational inheritance of *H3K27M^tet^*-induced defects from P0 to F1 generations, and MES-4 mediates the transgenerational inheritance of those defects in generations after F1.

### HERI-1 is required for both initiation and maintenance of TEI

Inhibition of HERI-1 or SET-32 can fully suppress the inheritance of *H3.3K27M^tet^*-induced defects from P0 to F1 generations **(Figures 2D and 5C)**. To test their dependencies on each other, we co-stained H3K23me3/H3K27me3 and HERI-1/H3K27me3 in HERI-1-or SET-32-depleted worms. *heri-1 RNAi* prevented the spread of H3K23me3 in *H3.3K27M^tet^* P0 **(Figure 6A)**, suggesting that the spread of H3K23me3 in *H3.3K27M^tet^* P0 requires HERI-1. Contrarily, *set-32 RNAi* did not influence the localization of HERI-1 to chromatin in *H3.3K27M^tet^* P0, revealing that HERI-1 can be targeted on chromatin in absence of H3K23me3, possibly through H3.3K27M **(Figure 3D)**. However, this chromatin localization was not maintained in the next generation in *set-32 RNAi* animals, and HERI-1 showed a diffuse wildtype distribution in the F1 **(Figure 6B)**, suggesting that the chromatin localization of HERI-1 in the F generations depends on the spread of H3K23me3 by SET-32. These results imply a read-and-write loop of HERI-1 and SET-32 to propagate H3K23me3, where HERI-1 recruits SET-32 upon the expression of *H3.3K27M^tet^* in the P0, and remains chromatin associated through binding to H3K23me3 in the *F1^subfertile^*, thus promoting the initiation of replicative TEI of the *H3.3K27M^tet^*-induced defects.

**Figure 6.**
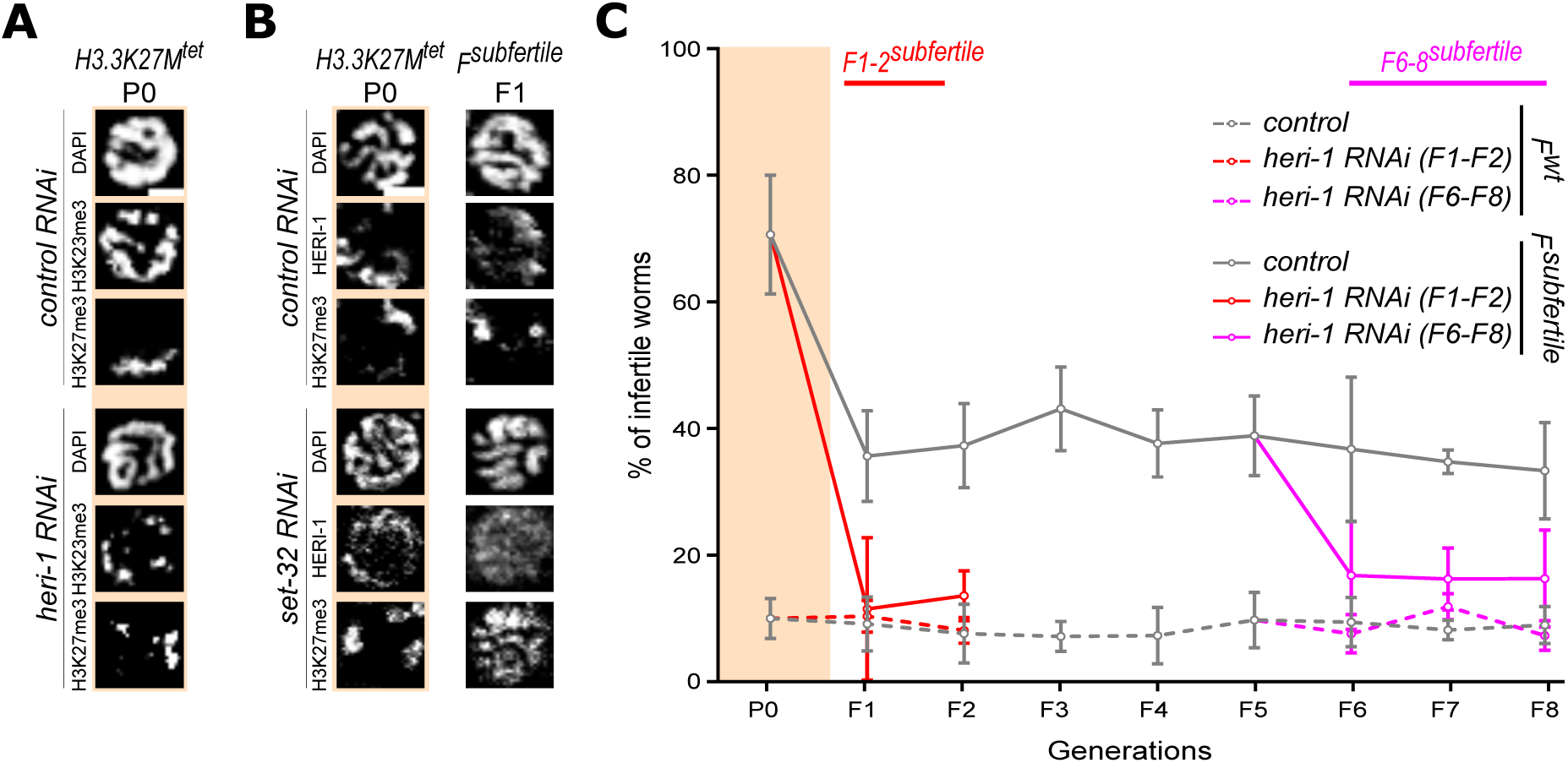
HERI-1 is required both for initiation and maintenance of TEI. **(A)** Representative images from immunostaining of H3K27me3 and H3K23me3 in late pachytene nuclei of *H3.3K27M^tet^*P0 with *control RNAi* or *heri-1 RNAi* treatment. **(B)** Representative images from immunostaining of H3K27me3 and FLAG-tagged HERI-1 in late pachytene nuclei of *H3.3K27M^tet^* P0-*F1^subfertile^* with *control RNAi* or *set-32 RNAi* treatment. Scale bars represent 2 µm in **(A, B)**. **(C)** Transgenerational infertility assay. Percentages of infertile worms in *P0-F8^wt^* (dashed lines) and *P0-F8^subfertile^* (solid lines) generations, with *control RNAi* (gray) or *heri-1 RNAi* treatment from F1 to F2 (red) or from F6 to F8 (fuchsia) at 25°C. Means and standard deviations of at least three biological replicates are shown.

In later *F^subfertile^* generations, inheritance of the *H3.3K27M^tet^*-induced fertility and chromatin defects can no longer be suppressed by *set-32 RNAi*, and the replicative TEI depends on the presence of MES-4 **(Figure 4C,D)**. Therefore, we wondered whether HERI-1 was only required for the initiation of TEI, or whether it also participated in the maintenance of TEI. Therefore, we targeted *heri-1* by RNAi in *F1-F2^subfertile^* and *F6-F8^subfertile^*. *heri-1 RNAi* fully suppressed the inherited fertility defects both in *F1-F2^subfertile^*and in *F6-F8^subfertile^* generations **(Figure 6C)**. HERI-1 is therefore involved in both the initiation and the maintenance of TEI, and bridges the intergenerational and transgenerational replicative EI of *H3.3K27M^tet^*-induced defects.

Based on these observations, we propose a mechanistic model for the TEI of altered H3K27me3 domains. In the initiation phase in the P0, *H3.3K27M^tet^*causes a depletion of autosomal H3K27me3 domains. CEC-6 recedes to the remaining H3K27me3 domains, while HERI-1 and SET-32 are recruited to chromatin and promote the replacement of H3K27me3 with H3K23me3, which reinforces chromatin localization of HERI-1. During the transition to the F generations, H3K36me3 replaces H3K23me3 and is required for the maintenance of the altered H3K27me3 patterns. During the maintenance phase, some germ cells can reconstruct wildtype H3K27me3 epialleles through CEC-6/PRC2. In other germ cells, HERI-1/MES-4 maintain the ectopic H3K36me3 epialleles, thus antagonizing the reconstruction of wildtype H3K27me3 domains **(Figure 7)**. This model explains the balance of replicative and reconstructive TEI mediated by the chromodomain proteins CEC-6 and HERI-1 and provides mechanistic insight into TEI of chromatin changes across generations.

**Figure 7.**
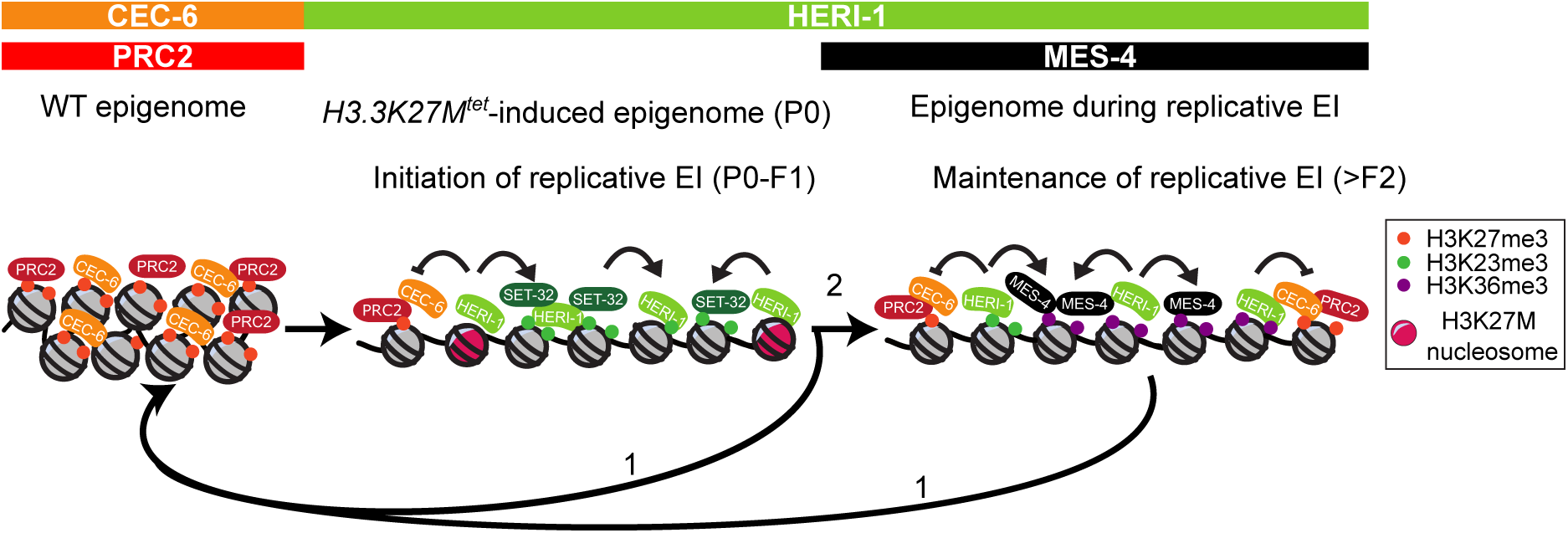
Model for the inheritance of *H3.3K27M^tet^*-induced chromatin defects. H3.3K27M incorporation into chromatin in the P0 induces the displacement of PRC2 and CEC-6, and the recruitment of HERI-1 to chromatin. After loss of the H3.3K27M protein, there are two inheritance paths of the epialleles to the subsequent generations: reconstructive EI (1) or replicative EI (2). For reconstructive EI (1), CEC-6 replaces HERI-1 and facilitates reloading of PRC2 to reconstruct wildtype H3K27me3 domains. For replicative EI (2), chromatin-bound HERI-1 recruits SET-32 to transiently spread H3K23me3 during the initiation phase (P0 to F1 transition). In the inheriting >*F2^subfertile^* generations H3K36me3 replaces H3K23me3, and MES-4 maintains the altered H3K36me3 domains in concert with HERI-1.

## Discussion

In this study, we demonstrate the TEI of altered H3K27me3 landscape for many generations in a wildtype *C. elegans* population. Many studies that have investigated EI have focused on the heritable (de)silencing of reporter transgenes by an acutely applied environmental stress^39^ or exogenously supplied sRNAs^17,40–44^. These studies have demonstrated that the effects on gene expression states can be passed on to unexposed offspring across several generations, requiring the RNAi machinery and involving chromatin modifications associated with gene silencing^17,27,45^. In parallel, studies describing the genetic removal of chromatin readers and writers provided insight into how chromatin marks are deposited and maintained^6,51–53^. Together, these studies clearly demonstrated the existence of TEI in *C. elegans*, but left open the questions whether TEI can rely on histone PTMs as inheriting factors, and how the changes in chromatin were mechanistically maintained across generations. Here, we revealed that an altered H3K27me3 landscape and the associated fertility defects can be transgenerationally heritable for many generations. Visualization of the H3K27me3 redistribution by IF allowed us to determine the persistence and penetrance of the inheritance in individual worms in different mutant background. We found that the TEI comprises two distinct steps: initiation, and maintenance **(Figure 7)**^15^. Both of these steps rely on a balance between the actions of the chromodomain proteins CEC-6 and HERI-1 **(Figures 2 and S5)**. In the initiation phase, the reduction of H3K27me3 results in the chromatin recruitment of HERI-1 **(Figure 3B)** and a feedforward loop with SET-32 to spread H3K23me3. During the maintenance of replicative EI, HERI-1 remains on chromatin, but H3K36me3 is deposited by MES-4 to replace H3K23me3. CEC-6 antagonizes the TEI and promotes the reconstruction of wildtype H3K27me3 patterns during both the initiation and the maintenance phases **(Figure 7)**. This competition between the replicative TEI and the reconstructive TEI resulted in a partial penetrance of the inherited fertility defects.

To experimentally select *subfertile* animals in each generation, we used an arbitrary cutoff of <50 offspring **(Figure S1C)**, resulting in an observation of 30-40% replicative TEI and 60-70% reconstructive TEI in each generation **(Figures 1B and 1H)**. Similarly, for quantifying germ cells with altered H3K27me3 landscapes, we only scored those that resembled cells expressing H3K27M in the P0. However, we observed a spectrum for both phenotypes, with a broad range of embryos laid **(Figure 1C)**, and intermediate stages of reconstruction of the wildtype H3K27me3 landscape **(Figure S2B).** The results resemble the low penetrance of epigenetically inherited phenotypes observed in previous studies^15,16,39,44–46,49–58^. The spectrum of germ cells with either replicated or reconstructed H3K27me3 distributions is observed within individuals **(Figure 1E)**, and not just within the population. The reconstruction occurred exclusively within autosomal genomic domains where H3K27me3 is present in wildtype worms (domains ii) **(Figure 1F)**. The observation of cells with partially reconstructed H3K27me3 landscapes **(Figure S2B)** suggests that this occurs gradually, but the ChIP experiments (where the reconstruction is averaged over many cells) suggest that there is no preferred order for domain reconstruction **(Figure 1F)**. We have previously shown that the global loss of H3K27me3 causes gene expression changes at many genes, but that de-silencing of the *kgb-1* gene is mainly responsible for the reduction in brood size^22^. The partial reconstruction of H3K27me3 domains in some germ cells therefore explains the imperfect correlation between chromatin defects and reduced brood sizes, as the latter depends on whether the *kgb-1* locus is targeted by the partial reconstruction. How specific regions are targeted by PRC2 for reconstruction is not clear. The amount of PRC2 is likely limiting this process, and our results show that CEC-6 enhances its function and pioneers the reconstruction of H3K27me3 domains. Recently, similar observations were made for sperm-inherited epialleles lacking H3K27me3, where the majority of loci regained H3K27me3 in the offspring, while few regions could not reconstruct H3K27me3, and instead maintained H3K36me3 marks^53,59^.

PRC2 is essential for development, as homozygous mutations in PRC2 result in defective lineage specification and body patterning in flies and mammals ^60–63^. Additionally, several monoallelic, recessive loss of function mutations in PRC2 (likely reducing the overall HMT activity) were associated with developmental disorders such as Weaver Syndrome, suggesting that not only the zygotic presence, but also the post-embryonic maintenance of H3K27me3 epialleles is essential in humans^64,65^. Therefore, defining the mechanisms underlying the maintenance of H3K27me3 is crucial for understanding cellular epigenetic memory in health and disease. In *C. elegans*, mutations in PRC2 result in a *mes* (maternal-effect sterile) phenotype due to the failure in the maintenance of silencing X-linked genes in the germ line^20,66^. This *mes* phenotype can be interpreted as an epigenetic phenotype, as loss of maternal PRC2 is only detrimental in the following generation, but the sterility prevents the analysis of future generations^21,67^. Here we take advantage of the inhibition of PRC2 by the transient expression of H3.3K27M^22^ to demonstrate that EI can persist beyond the following generation. This approach resembles the transient inhibition of PRC2 recently described in *Drosophila*, which leads to an irreversible derepression of genes linked to tumorigenesis^67^. Our results show that such a derepressed state can be heritable and under selection extend for many generations in *C. elegans*.

So far, no specific DNA sequence elements have been identified in *C. elegans* that are required for recruiting PRC2, and would thus be analogous to polycomb response elements (PREs) in *Drosophila*^68^ or CG islands in mammals^64^. Our observations suggest that instead, the de novo writing activity of PRC2 is relying on the chromodomain protein CEC-6 in *C. elegans*. The failure to (re-)initiate H3K27me3 domains explains why infertility phenotypes appear only after many generations in *cec-6* mutants, as replicative EI is still functional, but reconstructive EI is lost in absence of CEC-6. Similarly, the chromodomain protein HERI-1 is required for the de novo deposition of H3K23me3 and H3K36me3. Our findings are consistent with the analysis of other *C. elegans* chromodomain proteins that were found to have differential expression and localization patterns in the *C. elegans* germ line by interacting with distinct sets of histone PTMs and their writers^70,71^. CEC-6 and HERI-1 are homologs of CBX family proteins, which are members of the PRC1 complex in mammals^72–74^. There are homologs of the PRC1 complex in *C. elegans*, but they are not germline-expressed and likely do not contribute to the regulation of H3K27me3 deposition^75^. Recently, CBX2, a mammalian homolog of CEC-6, was proposed to demarcate H3K27me3 domains and recruit PRC2 to orchestrate cell differentiation^76^. This suggests that histone PTM reader proteins are required for regulating HMTs and controlling the de novo deposition and inheritance of chromatin landscapes, even though many HMTs also have histone PTM reader activity^77^.

The H3K27me3 and H3K36me3 domains are mutually exclusive in several model organisms^53,78,79^. We found that the switch from H3K27me3 domains to H3K36me3 domains required the transient deposition of H3K23me3 by SET-32. Maybe because of the transient nature of H3K23me3, point mutations at H3K23 or loss-of-function mutations of H3K23me3 writer or reader proteins were not associated with any obvious defects^80^. Consistently, mutants of *set-32* and *heri-1* are viable^15,58,81^. However, HERI-1 and SET-32 were previously identified in epigenetic suppressor screens^27,51,82,83^. SET-32 and/or HERI-1 were found to be required for the initiation of heritable epigenetic phenotypes such as an increased lifespan caused by the reduction of H3K4me3 levels or the silencing of a GFP reporter transgene^15,16,27,42,51^. In line with these previous studies, we found that HERI-1 and SET-32 do not have a suppressor effect on *H3.3K27M^tet^*P0, but are required for the initiation of replicative EI to the offspring. We propose that H3K23me3 is a transitory histone PTM required to initiate a heritable epigenetic state in response to induced changes in the epigenome.

The balance between reconstructive and replicative TEI by opposing histone PTMs in the germ line appears to be a general principle for heritable gene expression states, and may provide plasticity and epigenetic variation across generations^12,84^. Here, we have shown the long-term inheritance of an altered H3K27me3 landscape, which is maintained by a transient spreading of H3K23me3 followed by H3K36me3, and balanced by the opposing actions of CEC-6 and HERI-1. This inheritance of epigenome-based phenotypes provides a mutation-independent mechanism for adaptation to the environment.

## Limitations of the study

The study investigates transgenerational epigenetic inheritance over many generations, and probes the epigenome at every generation in a specific developmental window of the germline. This allows to determine what changes from one generation to the next, but does not allow to follow the dynamic processes that occur during development. Another limitation is that the stochastic nature of the epigenetic inheritance, and the relatively low penetrance of the phenotypes due to the selective pressure of (in)fertility, prevent us from generating large epigenetically homogenous populations that would be required for omics analyses in later generations.

## Methods

### *C. elegans* strains and culture

*C. elegans* strains were grown using standard conditions for maintenance at 20°C^85^. All experiments were performed at 25°C unless otherwise stated. Animals were maintained on either regular NGM (0.25% peptone, 0.3% NaCl, 1.5% agar, 1 mM CaCl_2_, 1 mM MgSO_4_, 25 mM KH_2_PO4 pH 6.0, 25 mM K_2_HPO4 pH 6.0 and 5 µg/mL cholesterol) seeded with *E. coli* OP50 or HT115, or on peptone rich NGM (2% peptone, 0.3% NaCl, 2% agar, 1 mM CaCl_2_, 1 mM MgSO_4_, 25 mM KH_2_PO4 pH 6.0, 25 mM K_2_HPO4 pH 6.0 and 5 µg/mL cholesterol) seeded with *E. coli* NA22 or HT115. N2 (Bristol strain) was used as wildtype. The generation of tetracycline-inducible transgenes expressing H3.3K27M was based on the sequence of the previously published *his-72(uge72)* allele containing a C-terminal OLLAS tag^22^. For most experiments, the transgene (referred to as *H3.3K27M^tet^*) was expressed from an extrachromosomal array obtained by microinjection of plasmids TC374 (containing the transgenic tetracycline-inducible driver)^24^ and pIO1 (containing the *his-72K27M::OLLAS* responder transgene). For native ChIP-seq experiments, to avoid mosaicism and low transmission rates of the *H3.3K27M^tet^* transgene, single copies of the transgenic tetracycline-inducible driver and responder transgenes were integrated into *ttTi5605* MosSCI^86^ and *jsTi1493* RMCE sites^87^, respectively **(Figure S1A)**. Regardless of the delivery, H3.3K27M always refers to a tetracycline-inducible transgenic copy of *his-72K27M::OLLAS* throughout the article. *H3.3K27M^tet^* induction was done on 1 ng/µl tetracycline-containing NGM plates seeded with HT115 unless otherwise indicated. All other strains were obtained either by CRISPR-Cas9 gene editing or genetic crossing. A list of strains used in this study is shown in **Table S3**.

### CRISPR-Cas9 gene editing

A 2xTEV::3xFLAG-tag was inserted at the C-terminus of the endogenous *cec-6* locus *(cec-6(ele23))* using a two-step CRISPR-Cas9 method^88^. This allele has a synthetic final intron containing a single loxP site. The chromodomain-dead *uge157ele23[cec-6(W55A,Y58A)]* and *uge158[heri-1(W62A,Y65A)]* point mutations were generated at the FLAG-tagged endogenous loci of *cec-6(ele23)* (in strain ALS611) and *heri-1* (in strain YY1481) using CRISPR-Cas9 as previously described in^89^. BsaJI and HaeII recognition sites were introduced as silent mutations into the *uge157ele23[cec-6(W55A,Y58A)]* and *uge158[heri-1(W62A,Y65A)]* alleles, respectively. Final verification of successful CRISPR-Cas9 edits was made by Sanger sequencing. sgRNAs were chosen based on the predicted quality score of CRISPRscan (https://www.crisprscan.org/)^90^. sgRNAs, repair templates, and PCR primers used to detect and sequence these edits are listed in **Table S4**.

### Plasmid generation

All generated plasmids including *H3.3K27M^tet^* transgenes and RNAi plasmids were cloned using standard Gibson assembly protocol with minor modifications^91^. Backbone plasmids were linearized by PCR or restriction digestion. PCR amplified inserts or hybridized oligos were ligated in 15 µl of Gibson mix (3.75% PEG-8000, 75 mM Tris-HCl (pH 7.5), 7.5 mM MgCl_2_, 0.15 mM dNTPs, 7.5 mM DTT, 0.75 mM NAD, 0.006 U/ul T5 exonuclease (NEB, M0363S), 0.025 U/µl Phusion DNA polymerase (NEB, M0530S), 4 U/µl Taq DNA ligase (NEB, M0208S)). The molar ratio of insert to backbone was 3:1. The mixture, 20 µl in total, was incubated at 50°C for 1 hour. The final check on candidate clones was done by Sanger or whole plasmid sequencing. **Table S5** contains the names of plasmids used in the study.

### RNAi treatment

The standard RNAi by feeding protocol^92^ was used. Briefly, overnight grown RNAi pre-cultures in LB medium containing ampicillin at 37°C were diluted 10x in the same medium and incubated for 3-4 hours at 37°C (until OD values reach up to 0.6-0.8), followed by 4 mM IPTG induction for another 4 hours at 37°C. The cultures were 2 times concentrated before seeding onto freshly prepared RNAi NGM (-/+ 1 ng/µl tetracycline, 1 mM IPTG, and 25 µg/ml carbenicillin) or peptone rich RNAi NGM (-/+ 30 ng/µl tetracycline, 1 mM IPTG, and 25 µg/ml carbenicillin). *cec-6, heri-1, cec-1 RNAi* clones were obtained from the Ahringer library^93^ while *mes-4 and set-32 RNAi* clones were generated by Gibson assembly **(Table S3)**.

### Transgenerational infertility assay

10-12 P-1 animals (parents of P0 generation) were selected based on the presence or absence of the extrachromosomal *H3.3K27M^tet^* array and placed at 25°C when they were at early L4 larval stage **(Figure 1A)**. From these P-1 plates, >50 L3/early L4 animals (before any phenotypes were apparent) of the P0 generation were singled. After 3-4 days, these P0 animals were scored based on the presence of gonad and brood size under a dissecting microscope (Leica S9E stereo microscope) into categories as in **Figure S1C**. *Fertile* animals lay more than 50 viable embryos. *No gonad* animals do not produce any offspring, while *subfertile* worms lay less than 50 viable embryos. For each subsequent F generation, >50 offspring of *fertile* or *subfertile* animals from the previous generation were randomly selected at L3/L4 stage and singled out for the analysis of phenotypes in the *F^wt^*, *F^subfertile^* or *F^fertile^* lineages. Fertility level comparisons were performed in at least three biological replicates, and the raw data are in **Tables S4** and **S5**. Statistical tests applied for each experiment were specified in the corresponding figure legends.

### Immunofluorescence staining and image analysis

Worm gonads were dissected in 4 µl of anesthetizing DB buffer (1.6% sucrose, 73 mM HEPES pH 6.9, 40 mM NaCl, 5 mM KCl, 2 mM MgCl_2_,10 mM EGTA pH 7.5, 0.1% NaN_3_) on 10 well microscope slides (Thermo Fisher Scientific) with a 21G needle (BD Microlance). Dissected gonads were transferred to poly-L-lysine-coated (0.1% (w/v); Sigma-Aldrich) 3-square-well microscope slides (Thermo Fisher Scientific). Slides were placed on a dry-ice cold metal plate for at least 15 min. Samples were fixed in methanol or ethanol with acetone **(Table S6)**. The fixatives were stored at -20°C and re-used for a maximum of 10 experiments. Slides were washed three times in PBS for 5 min and incubated with 1:300 anti-ollas (Novus Biologicals NBP1-06713), 1:300 anti-H3K27me2/3 (Active motif 39535), 1:50 anti-FLAG (Sigma F7425), 1:600 anti-H3K23me3 (Active motif 61499) or 1:2000 anti-H3K36me3 (Abcam ab9050) antibodies overnight at 4°C in a humid chamber. Samples were washed in PBS-Tween 20 three times for 5 min and incubated with 1:1000 times diluted Cy3-(Jackson ImmunoResearch) or Alexa488 (Thermo Fisher Scientific) secondary antibodies for 1.5 hours at room temperature in the dark. Samples were washed with PBS-Tween 20 three times for 5 min, stained with 2 µg/ml DAPI for 15 min, and mounted with VECTASHIELD Antifade Mounting Medium (H1200). 2-D and confocal images were obtained using a Leica DM5000B with 63x oil objectives (NA = 1.25) and a Leica LSM800 confocal microscope with 63x oil objectives (NA = 1.40), respectively. Representative confocal images show maximum projections of 0.1 μm Z-sections of the entire pachytene nuclei.

For quantifying the extent of co-localization of chromodomain proteins or histone marks on the chromatin of stained pachytene nuclei, MMMEA macro and EZcolocalization plugin in Fiji were used^94,95^. The maximum Z projections for each nuclei were stacked based on DAPI signal. Graphs were generated based on the user-defined parameters and the obtained measurements using a custom R script. The percentage of pachytene cells showing altered H3K27me3, H3K36me3 and H3K23me3 patterns per gonad were scored manually using FiJi^95^.

### Native ChIP-seq

The native ChIP protocol was adapted from^96^. L1 animals synchronized by sodium hypochlorite treatment were grown on 14-cm peptone-rich NGM plates. P-1 and P0 generations containing single copy integrated *H3.3K27M^tet^*, wildtype controls, and animals with respective genetic background or RNAi treatment were grown on 30 ng/µl tetracycline-containing plates seeded with HT115, whereas the inheriting F generations were grown on tetracycline-free plates seeded with NA22, or HT115 expressing for strains with RNAi treatment. Worms were harvested as young adults, washed in M9 (22 mM KH_2_PO_4_, 42 mM K_2_HPO_4_, 85 mM NaCl, 1 mM MgSO_4_) at least 6 times, and resuspended in buffer A (15 mM Tris pH = 7.5, 2 mM MgCl_2_, 340 mM sucrose, 0.2 mM spermine, 0.5 mM spermidine, 0.5 mM PMSF) supplemented with 0.5% NP-40 and 0.1% Triton X-100. Total nuclei, which are enriched for germ-cell nuclei based on morphologic analysis, were obtained by light grinding of the worms under liquid nitrogen followed by douncing, low-speed centrifugation and washes. 150 μl of chromatin was digested into nucleosome ladders (mono, di, tri…) using 5 μl of MNase (NEB M0247S) in 850 μl of 5 mM CaCl_2_ containing buffer at 37°C by shaking for 1, 3, 5, 8, 10, 12, 15 and 20 min. At each time point, ∼120 μl of the sample was removed and the MNase inhibited by the addition of 50 mM EDTA pH 8. The samples from the different timepoints were pooled to obtain nucleosomal ladders. Following digestion, chromatin was extracted, solubilized, and pre-cleared with empty beads. The soluble chromatin fraction was incubated with 1:250 primary antibodies against H3K27me2/me3 (Active Motif 39535) overnight at 4°C. Antibody-bound chromatin was captured with magnetic beads for 1 hour at 4°C (50 μl 1:1 mixture of Dynabeads Protein A and Dynabeads Protein G, Invitrogen). DNA from both input and ChIP samples was extracted using phenol/chloroform. Libraries were prepared using NEBNext® Ultra™ II DNA Library Prep with sample AMPure XP magnetic purification beads (Beckman Coulter, A63881) using 100 ng of input DNA. Libraries were sequenced using the TruSeq SBS HS v3 chemistry on an Illumina HiSeq 2500 or NovaSeq 6000 sequencer and demultiplexed using dedicated 6-mer Illumina TruSeq indexes.

### ChIP-seq data analysis

Paired-end reads (length=150 bp) were mapped to the *C. elegans* reference genome WBcel235 (*ce11*) with Bowtie2 (default parameters, producing BAM format files) using Galaxy^97^. Poor-quality reads and reads from blacklisted regions^98^ were removed using the BAM filter tool. All files were normalized to the same number of reads. Read counts in the IP files were normalized to read counts in the input files by division followed by log2 transformation. Genome browser tracks were inspected for each individual replicate. Aligned reads (BAM files) of biological replicates of ChIP-seq experiments that passed the quality control (minimum of three) were merged, and the downstream pipeline was performed again using the merged BAM files. Processed data files are bedgraphs containing normalized ChIP results. Bin sizes used in the analysis were 200 bp and each bin was smoothed to 2 up- and 2 downstream bins. Genome browser images were obtained using IGV 2.12^99^. H3K27me3 occupancy domains were defined as in^22^, and the occupancies determined and plotted using a custom R script.

### Purification of recombinant proteins from bacteria

cDNA sequences encoding N-terminally 10xHis-MBP-TEV tagged CEC-6 (aa2-120) and HERI-1 (aa2-127) were cloned into pelB bacterial expression plasmids by Gibson assembly. Domain borders were identified based on Alphafold2 predictions^31,32^. The recombinant proteins were overexpressed in *E. coli* BL21, grown in 2xYT media at 37°C for ∼4 hours (until an OD of 0.6) followed by 1 mM IPTG induction at 18°C for 6 hours before cell harvest. All subsequent steps were performed at 4 °C. Induced cells were harvested by centrifugation and resuspended in lysis buffer (20 mM HEPES pH 8.0, 300 mM NaCl, 10% glycerol, 5 U/ml supernuclease (Novagen, SSNP01), protease inhibitor cocktail tablets (complete EDTA-free; Roche Diagnostics). Cells were lysed using an Emulsiflex system (AVESTIN) and cleared by centrifugation at 32000g for 1 hour at 4°C. The soluble fraction was loaded onto a HisTrap HP column (Cytivia) on an AKTA-HPLC purifier (GE Healthcare). Samples were eluted by applying an imidazole gradient reaching from 15 mM imidazole to 500 mM imidazole final concentration in the lysis buffer. Peak fractions were pooled, and the NaCl concentration was reduced to 100 mM before loading onto a heparin affinity column (Cytivia). Protein samples were eluted by applying a salt gradient ranging from 100 mM NaCl to 1 M NaCl final concentration in the buffer solution. Peak fractions were pooled and subsequently loaded on a Superdex 75 Increase 10/300 GL column (Cytivia). The pure proteins obtained from size-exclusion chromatography were concentrated to 0.5-1.5 mg/ml and snap-frozen and stored at -80°C **(Figure S6A)**.

### In vitro histone peptide pulldown assay

N-terminally biotinylated 20 aa peptides **(Table S7)** of histone H3 (unmodified, K27M mutation or with trimethylated lysines at different positions) were chemically synthesized (Genscript). 60 µl histone peptides (7.5 µM) were coupled to 20 µl Dynabeads M-280 streptavidin beads (Invitrogen) in binding buffer (20 mM HEPES pH 8, 600 mM NaCl, 5% glycerol, 0.1% Triton X-100) rotating for 1 hour at 4°C. 40 µl of 20 µM purified recombinant CEC-6 and HERI-1 MBP fusion proteins were incubated with the histone peptide-coupled beads slurry in binding buffer for 1 hour at 4°C on a rotator. After washing five times with the binding buffer, bound proteins were released from the beads by boiling in SDS Laemmli buffer for 5 min and resolved by SDS-PAGE gel electrophoresis followed by QuickBlue Protein stain (LubioScience). Band intensities were quantified using FiJi^95^. After subtracting the background signal value (signal from no peptide treatment), band intensity values of each pull-down sample were normalized by division with the input band intensity value.

### Sequence and structure comparison of CEC-6 and HERI-1

The annotated chromodomain sequence of CEC-6^26^ was BLASTed to the *C. elegans* proteome in Uniprot (https://www.uniprot.org/blast). One of the top hits was the HERI-1 chromodomain. The structures of the two chromodomains and the Pc protein from *D. melanogaster* were predicted using Alphafold2^100^ with default parameters. Predicted structures were superimposed in ChimeraX^101^.

### Data representation and statistical analysis

All the data analysis and statistical significance testing was done using the GraphPad Prism 9.5 software package (GraphPad) and R studio. The figures were assembled in Inkscape.

## Data and code availability

All relevant data supporting the findings of this study are available within the article and its Supplementary Information files. Further information and requests for resources and reagents should be directed to and will be fulfilled by the lead contact, Florian Steiner. Raw data underlying the figures will be deposited at YARETA. Sequencing data has been deposited at the Gene Expression Omnibus (GEO) under accession number GSE270029.

## Reporting summary

Further information on research design for this article is available as a Supplementary Information file.

## Acknowledgments

We are grateful to the present and past members of the Steiner laboratory, Michaela Dohnálková and Monica Gotta for their comments and suggestions. We thank the Bioimaging Center of the Faculty of Sciences and iGE3 Genomics Platform at the University of Geneva. Some strains were provided by the CGC, which is funded by NIH Office of Research Infrastructure Programs (P40 OD010440). We thank WormBase and the worm community, especially Scott Kennedy, Sam Guoping Gu, Shouhong Guang, and Michael L. Nonet for kindly providing strains and reagents. The work was financially supported by the Swiss National Science Foundation (Grants 310030_197762 and 31003A_175606 to FAS), and funding from the Republic and Canton of Geneva.

## Author contributions

IO and FAS conceptualized the ideas and designed experiments. IO performed the majority of the experiments. AH contributed to the purification of the chromodomains, and CL generated ALS611 *(cec-6(ele23))* strain. JMW and KD discovered the epigenetic heritability of infertility. AS, AB and FAS supervised the work and acquired funding. IO and FAS wrote the manuscript with input from all the authors.

---

**Figure S1.**
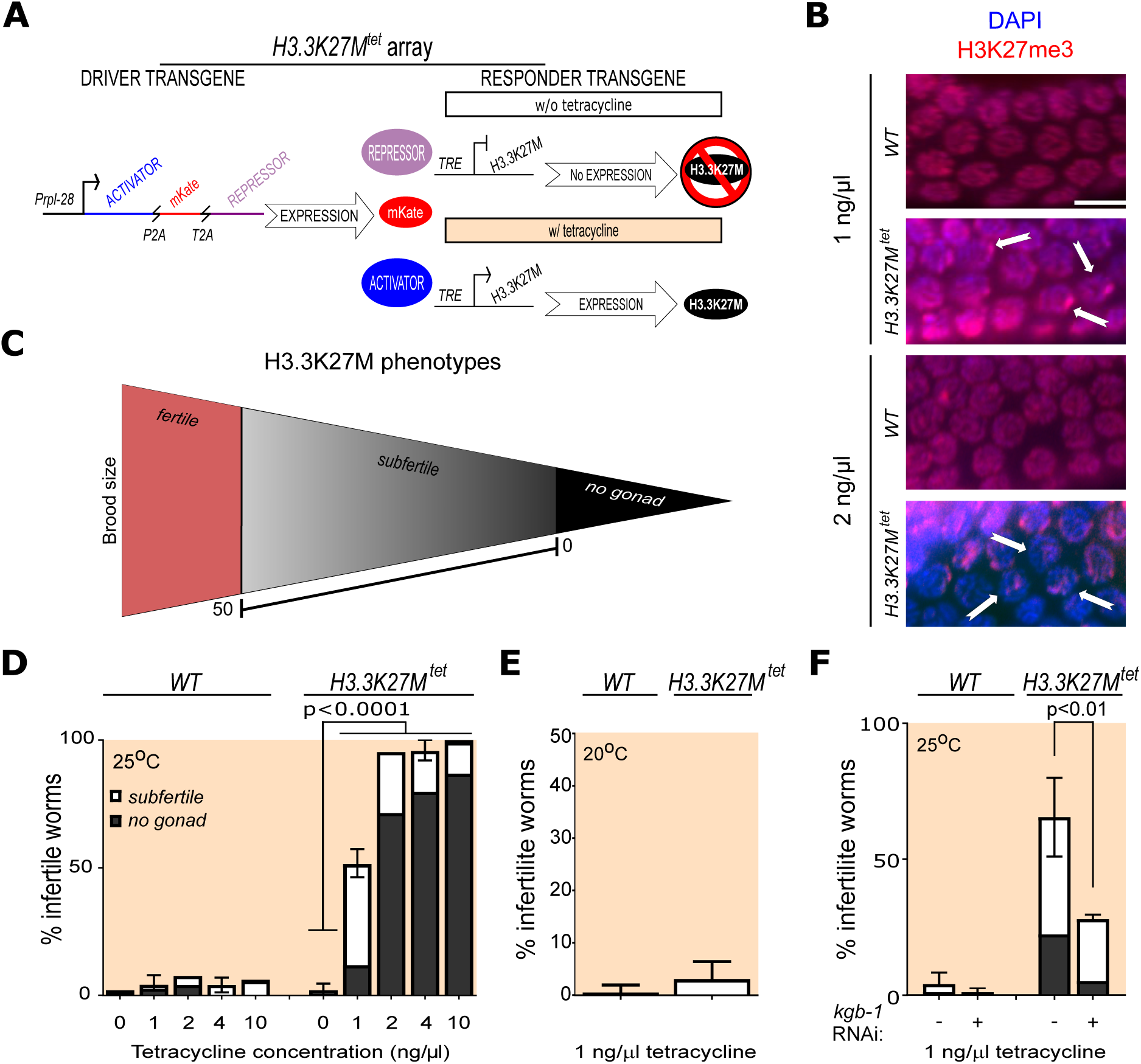
Inducible expression of *H3.3K27M^tet^* initiates changes in H3K27me3 distribution and fertility defects. **(A)** Schematic representation of tetracycline-inducible expression of *H3.3K27M^tet^*. *H3.3K27M^tet^*consists of two co-injected transgenes: i) the driver transgene constitutively co-expressing the activator (rTetR+QFAD), mKate (fluorescent marker), and the repressor (TetR+*C. elegans pie-1* repressor domain) and ii) the responder transgene consisting of regulatory TRE (*7xTetO::Ppes-10*) controlling the coding sequence for OLLAS-tagged H3.3K27M. ACTIVATOR, mKate and REPRESSOR are separated by self-cleaving P2A and T2A peptides. Upon tetracycline exposure, ACTIVATOR replaces REPRESSOR binding to the TRE and drives the expression of *H3.3K27M^tet^*. For details, see methods and^24^. **(B)** Representative images from immunostaining of H3K27me3 in wildtype and *H3.3K27M^tet^* pachytene nuclei from animals treated with 1 ng/µl or 2 ng/µl tetracycline. Examples of nuclei with an altered H3K27me3 landscape are highlighted with arrows. Scale bar represents 5 µm. **(C)** Cartoon showing the spectrum of fertility defects of *H3.3K27M^tet^.* Subfertility was arbitrarily defined as having a brood size of less than 50 offspring. **(D-F)** Quantification of infertility of wildtype and *H3.3K27M^tet^* animals. **(D)** Worms were treated with 1 ng/µl, 2 ng/µl, 4 ng/µl and 10 ng/µl tetracycline at 25°C. **(E)** Worms were treated with 1 ng/µl tetracycline at 20°C. **(F)** Worms were treated simultaneously with 1 ng/µl tetracycline, and *control RNAi* or *kgb-1 RNAi* at 25°C. Means and standard deviations of at least three biological replicates are shown. P-values are from 2-way ANOVA with Sidak’s multiple comparison tests.

**Figure S2.**
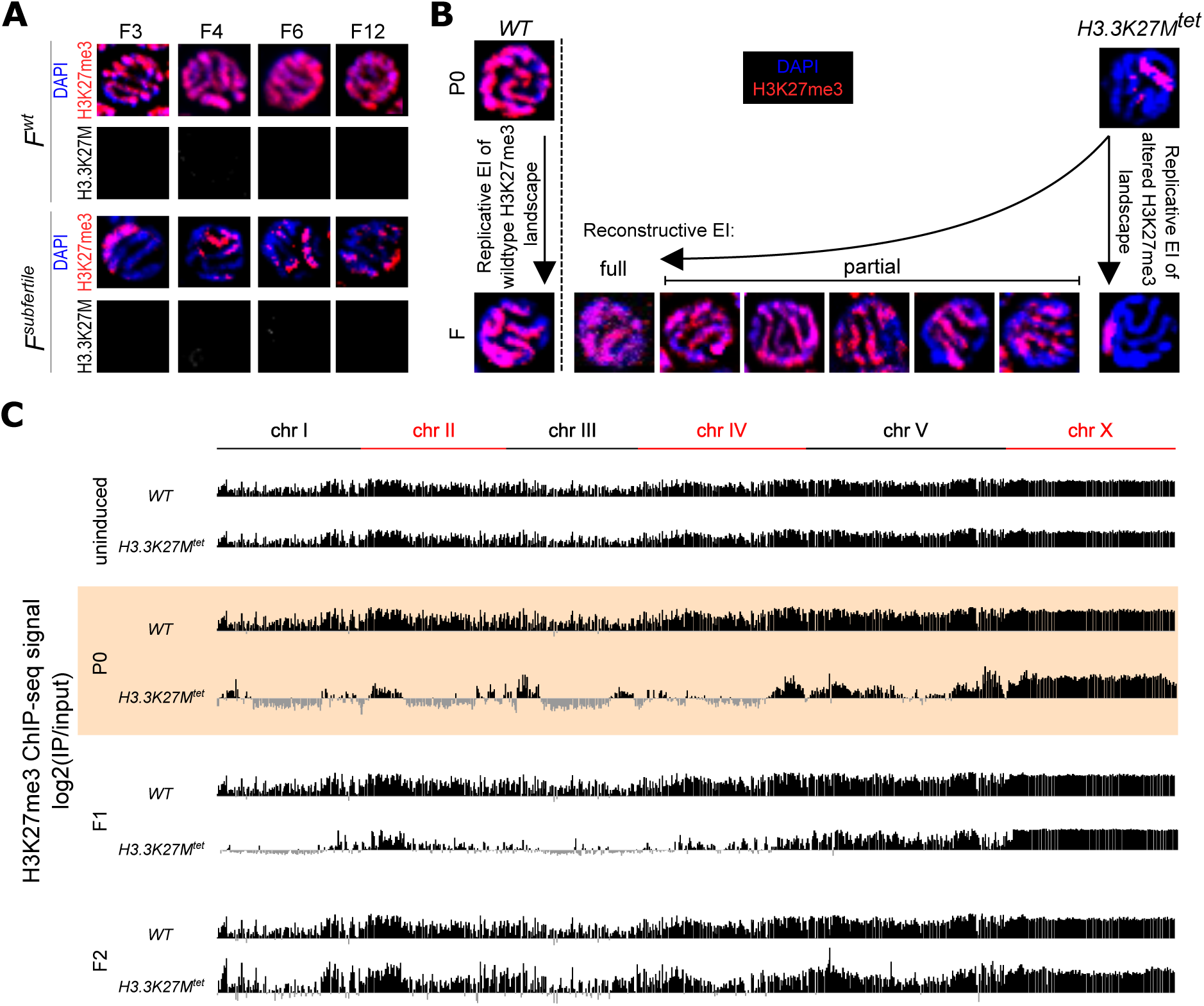
*H3.3K27M^tet^*-induced changes in the H3K27me3 landscape are subject to replicative and reconstructive EI. **(A)** Representative images from immunostaining of H3K27me3 and OLLAS-tagged H3.3K27M in late pachytene cells in generations F3, F4, F6 and F12 of *F^wt^* and *F^subfertile^*. Generations P0, F1, F2, F8 and F15 are shown in Figure 1D. **(B)** Spectrum of the H3K27me3 landscapes in late pachytene cells observed during reconstructive EI, with replicative EI of wildtype and *H3.3K27M^tet^*-induced altered landscapes shown for comparison. For quantifications of % nuclei with altered H3K27me3 landscapes per gonad in Figure 1E, only complete replicative EI was considered as “altered”, whereas full and partial reconstructive EI was considered as “wildtype”. **(C)** Genome browser views of genome-wide H3K27me3 levels as determined by H3K27me3 ChIP-seq experiments from uninduced, tetracycline-induced P0, and F1 and F2 generations in wildtype (WT) and *H3.3K27M^tet^*strains at 25°C. Each ChIP-seq track is the average log_2_ ratio (IP/input) from three independent biological replicates. Individual chromosomes are represented by black and red lines above the tracks.

**Figure S3.**
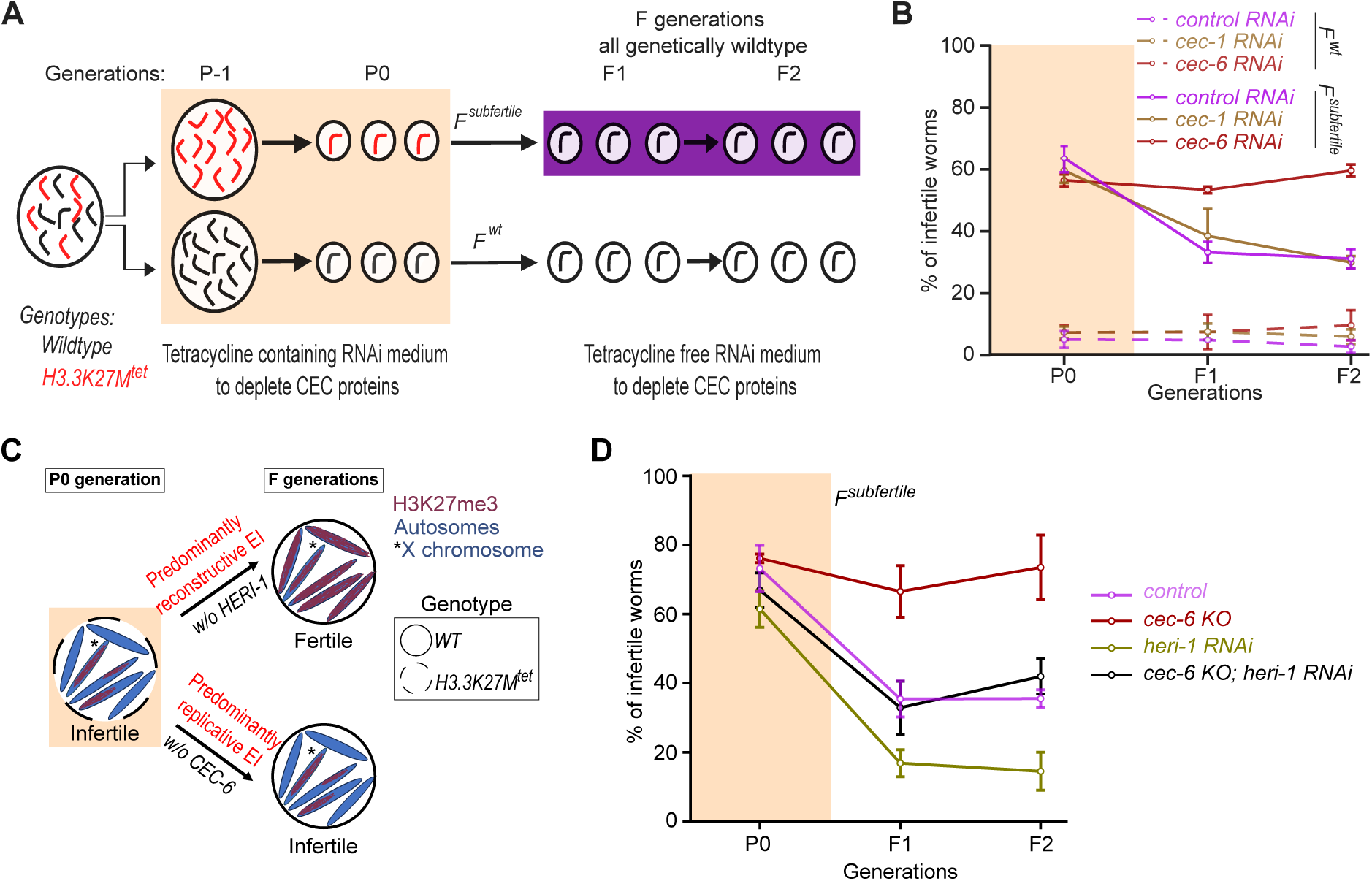
CEC-6 and HERI-1 balance the inheritance of the H3K27me3 landscape. **(A)** Schematic representation of RNAi depletion of chromodomain proteins during the TEI assay, similar to Figure 1A. Wildtype (black) and *H3.3K27M^tet^* (red) animals were exposed to tetracycline and RNAi during P-1 and P0 generations (beige background), and their *F^wt^* (white background) and *F^subfertile^* (purple background) descendants were exposed to RNAi at 25°C on plates without tetracycline. **(B)** Percentages of infertile worms in *P0-F2^wt^* (dashed lines) and *P0-F2^subfertile^*(solid lines) generations upon *control RNAi*, *cec-1 RNAi* or *cec-6 RNAi* at 25°C. **(C)** Cartoon of pachytene nuclei summarizing replicative and reconstructive EI of *H3.3K27M^tet^-* induced defects from P0 to inheriting F generations in absence of CEC-6 or HERI-1, respectively. CEC-6 and HERI-1 do not influence the *H3.3K27M^tet^*-induced chromatin changes and associated fertility defects in the P0 generation, but promote (w/o CEC-6) or oppose (w/o HERI-1) the inheritance of these defects. The *H3.3K27M^tet^* expressing pachytene nucleus is shown with a dashed border, and genetically wildtype nuclei are shown with solid borders. Chromosomes are shown in blue, and the X chromosome is marked by an asterisk. The H3K27me3 distribution is shown in magenta. **(D)** Percentages of infertile worms in *P0-F2^subfertile^* generations upon *control RNAi*, *cec-6 KO*, *heri-1 RNAi*, or *cec-6 KO* and *heri-1 RNAi* at 25°C. **(B, D)** Means and standard deviations of at least three biological replicates are shown.

**Figure S4.**
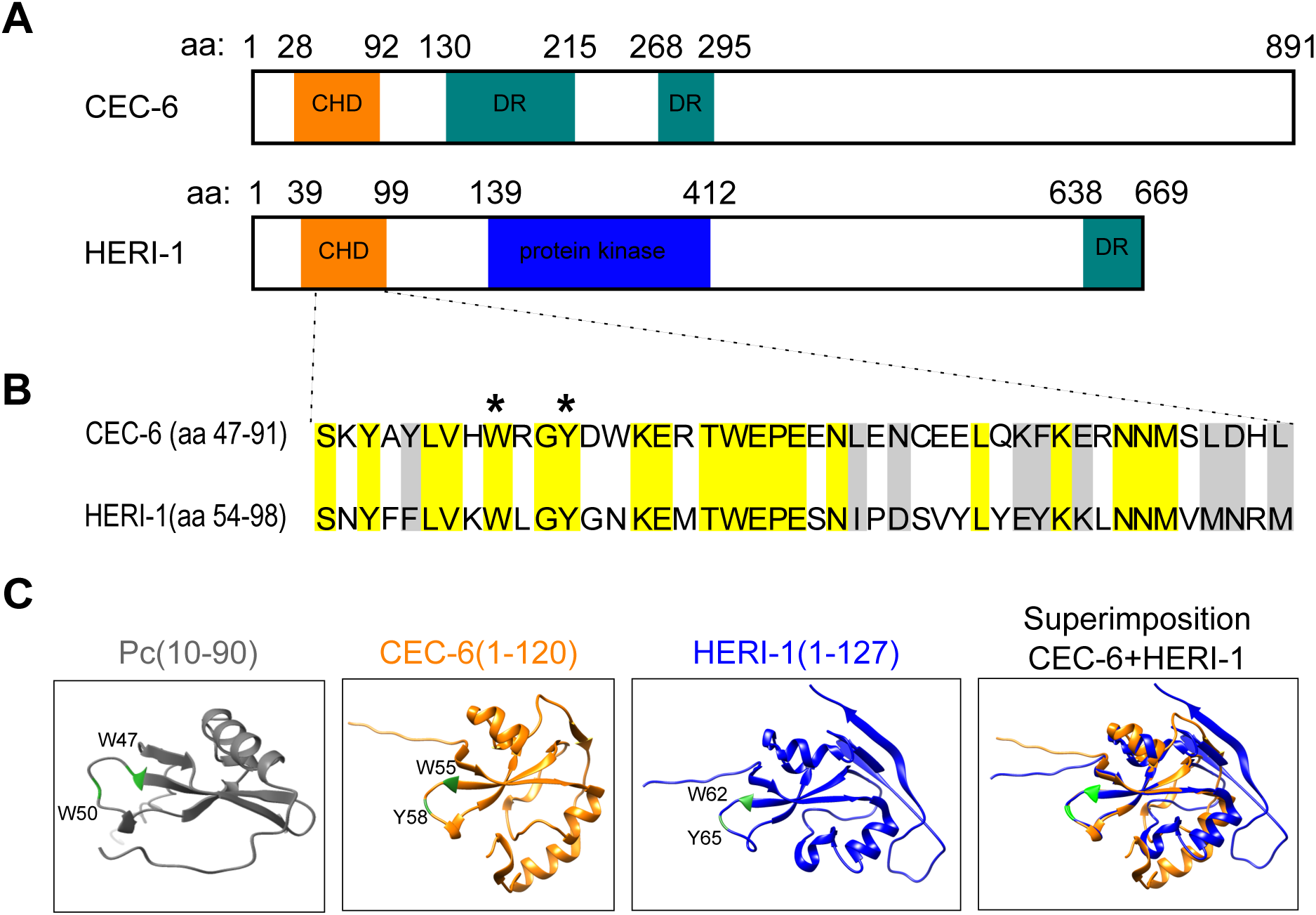
The chromodomains of CEC-6 and HERI-1 are similar and resemble the chromodomain of *Polycomb* (Pc) in *Drosophila melanogaster*. **(A)** Schematic representation of full-length CEC-6 and HERI-1 proteins with annotated chromodomains (CHD, orange), disordered regions (DR, green) and a putative protein kinase domain (HERI-1 only, blue). **(B)** Amino acid sequence comparison of the chromodomains of CEC-6 (aa47-91) and HERI-1 (aa54-98). Identical amino acids are highlighted in yellow, amino acids with similar properties are highlighted in gray. Asterisks indicate aromatic amino acids essential for methyl lysine binding (CEC-6W55 and Y58, HERI-1W62 and Y65). **(C)** Alphafold2-predicted structures of *Drosophila melanogaster* Pc(aa10-90) (gray), CEC-6(aa1-120) (orange) and HERI-1(aa1-127) (blue). A superimposition of CEC-6(aa1-120) and HERI-1(aa1-127) is shown on the right. The aromatic amino acid residues essential for methyl lysine binding marked with asterisks in (**B)** are highlighted in green.

**Figure S5.**
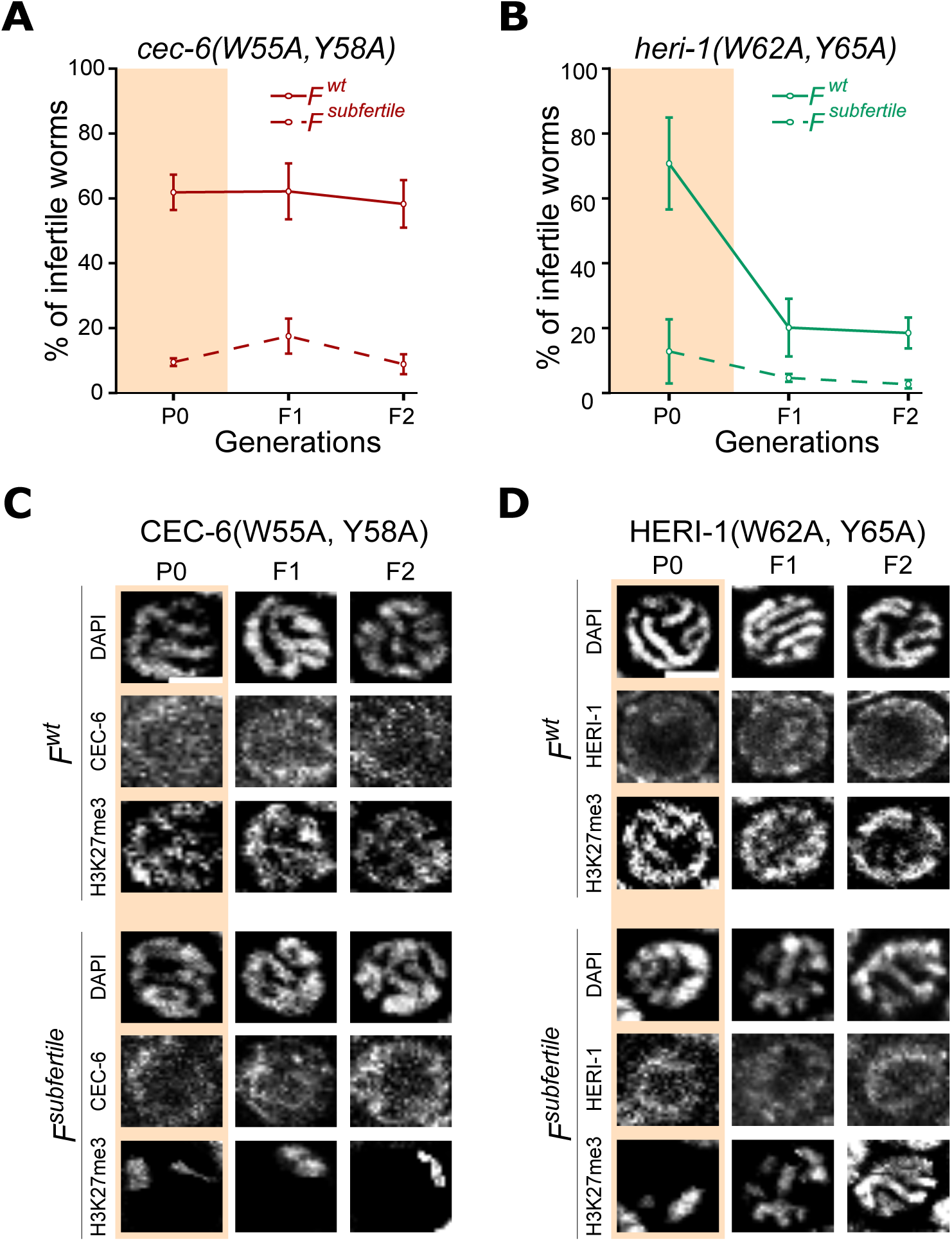
CEC-6(W55A, Y58A) and HERI-1(W62A, Y65A) point mutations abolish their chromatin localization and their roles in replicative and reconstructive EI of *H3.3K27M^tet^*-induced defects. **(A and B)** Percentages of infertile worms in *P0-F2^wt^* (dashed lines) and *P0-F2^subfertile^* (solid lines) generations in **(A)** *cec-6(W55A, Y58A)* mutants, or **(B)** *heri-1(W62A, Y65A)* mutants. Means and standard deviations of at least three biological replicates are shown. **(C and D)** Representative images from immunostaining of H3K27me3 and FLAG-tagged chromodomain proteins in late pachytene nuclei in *P0-F2^wt^*and *P0-F2^subfertile^* generations in **(C)** *cec-6(W55A, Y58A)* mutants or **(D)** *heri-1(W62A, Y65A)* mutants. Scale bars represent 4 µm.

**Figure S6.**
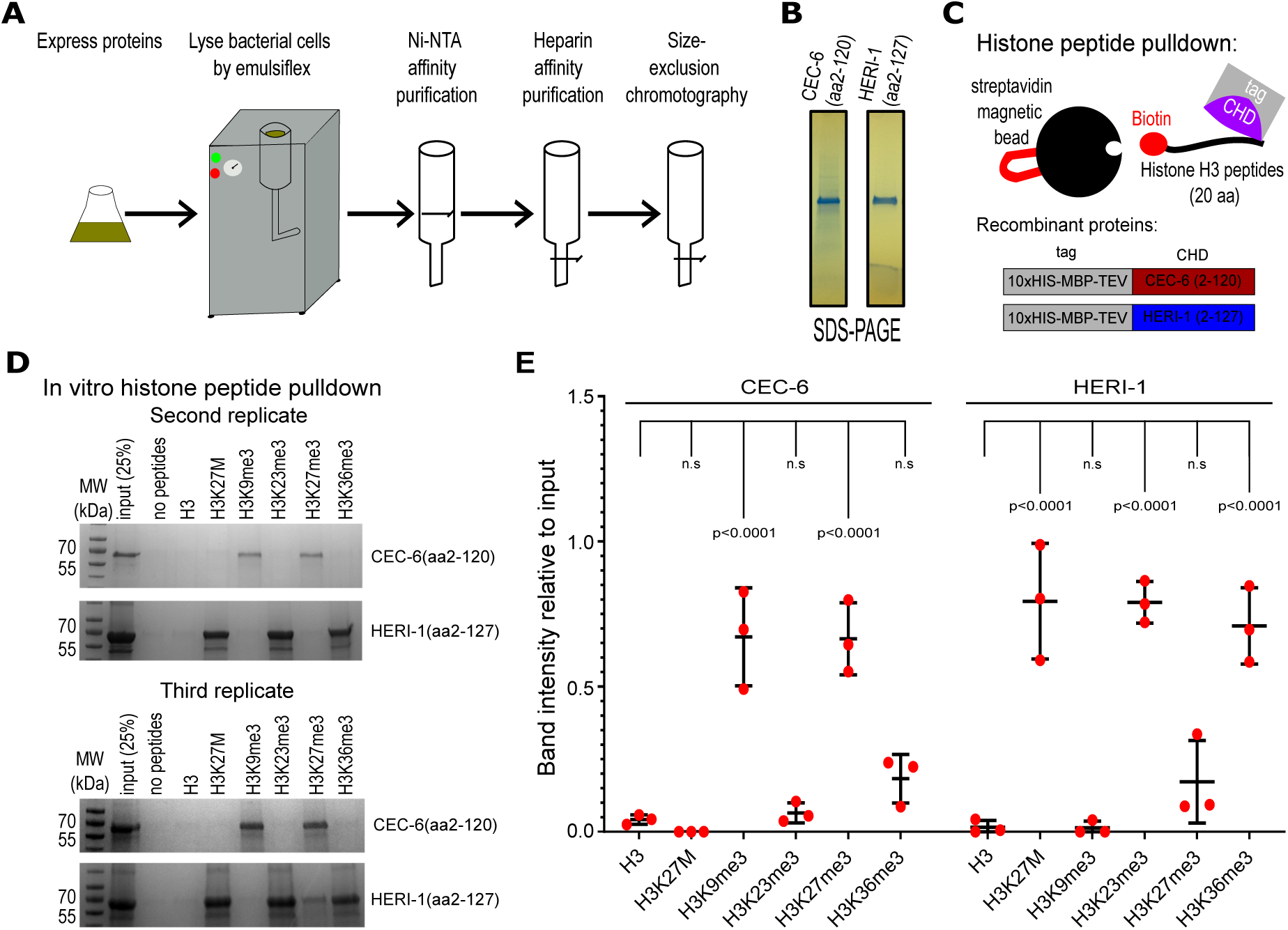
CEC-6 and HERI-1 chromodomains bind to distinct histone PTMs in vitro. **(A)** Purification scheme of recombinant 10xHis-MBP-tagged chromodomains of CEC-6(aa2-120) and HERI-1(aa2-127) from *E. coli*. Cells expressing the proteins were lysed, and lysates passed over a Ni-NTA affinity column, then a heparin affinity column, and finally a size-exclusion column. **(B)** Coomassie staining of purified recombinant 10xHis-MBP-tagged CEC-6(aa2-120) and HERI-1(aa2-127) proteins after SDS-PAGE. **(C)** Cartoon of in vitro histone peptide pulldown of 10xHis-MBP-tagged CEC-6(aa2-120) and HERI-1(aa2-127). The recombinant proteins were mixed with magnetic streptavidin beads bound to biotin-labeled histone H3 peptides with the corresponding PTMs and co-precipitated using a magnet. **(D)** Coomassie stainings of second and third replicates of in vitro histone peptide pulldown of CEC-6(aa2-120) and HERI-1(aa2-127) as in Figure 3D. **(E)** Quantifications of relative band intensities of CEC-6(aa2-120) and HERI-1(aa2-127) pulled-down with H3 peptides carrying different PTMs in gels shown in Figures 3D **and S6D**. Means and standard deviations from the three biological replicates are shown. P-values are from 2-way ANOVA with Tukey’s post hoc test in comparison to the unmodified H3 peptide. n.s. stands for not significant.

**Figure S7.**
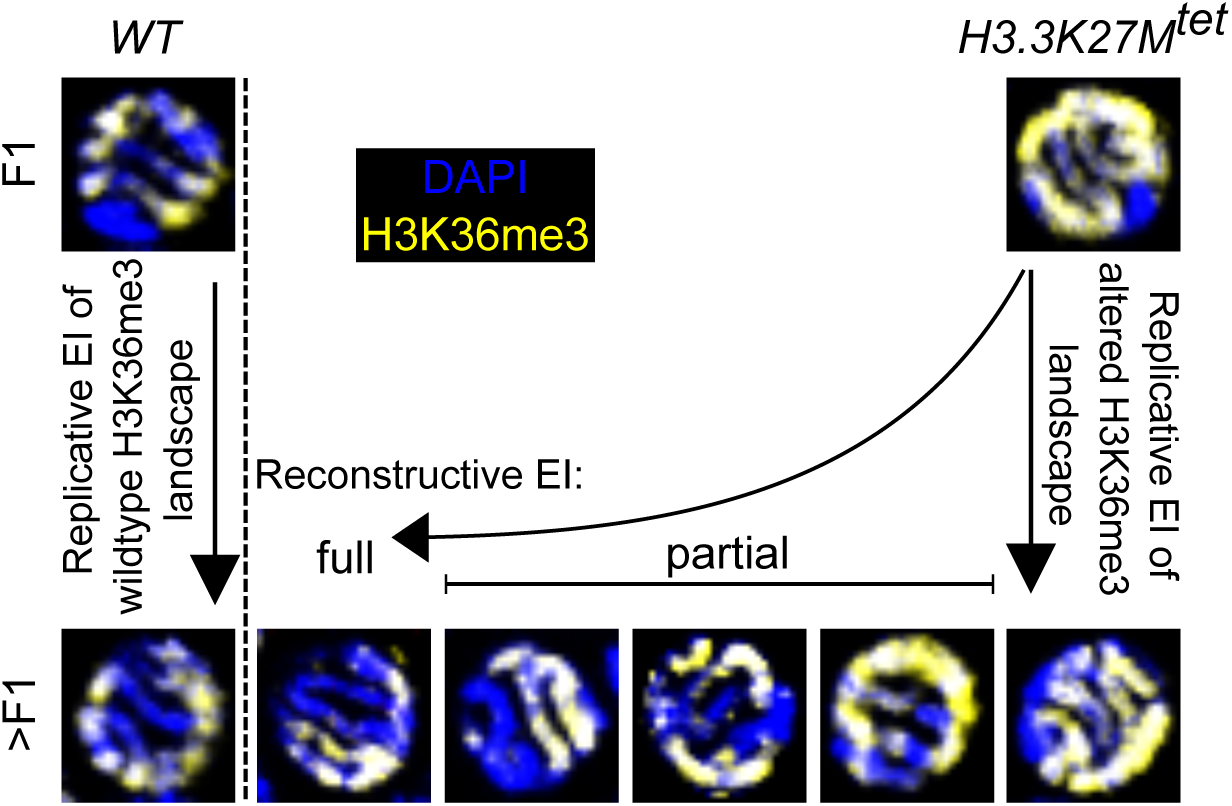
Altered H3K36me3 landscapes are subject to replicative and reconstructive EI in Fsubfertile. Spectrum of the H3K36me3 landscapes in late pachytene cells observed during reconstructive EI, with replicative EI of wildtype and altered H3K36me3 landscapes shown for comparison. For quantifications of % nuclei with altered H3K36me3 landscapes per gonad in Figure 4B, only complete replicative EI was considered as “altered”, whereas full and partial reconstructive EI was considered as “wildtype”.

**Figure S8.**
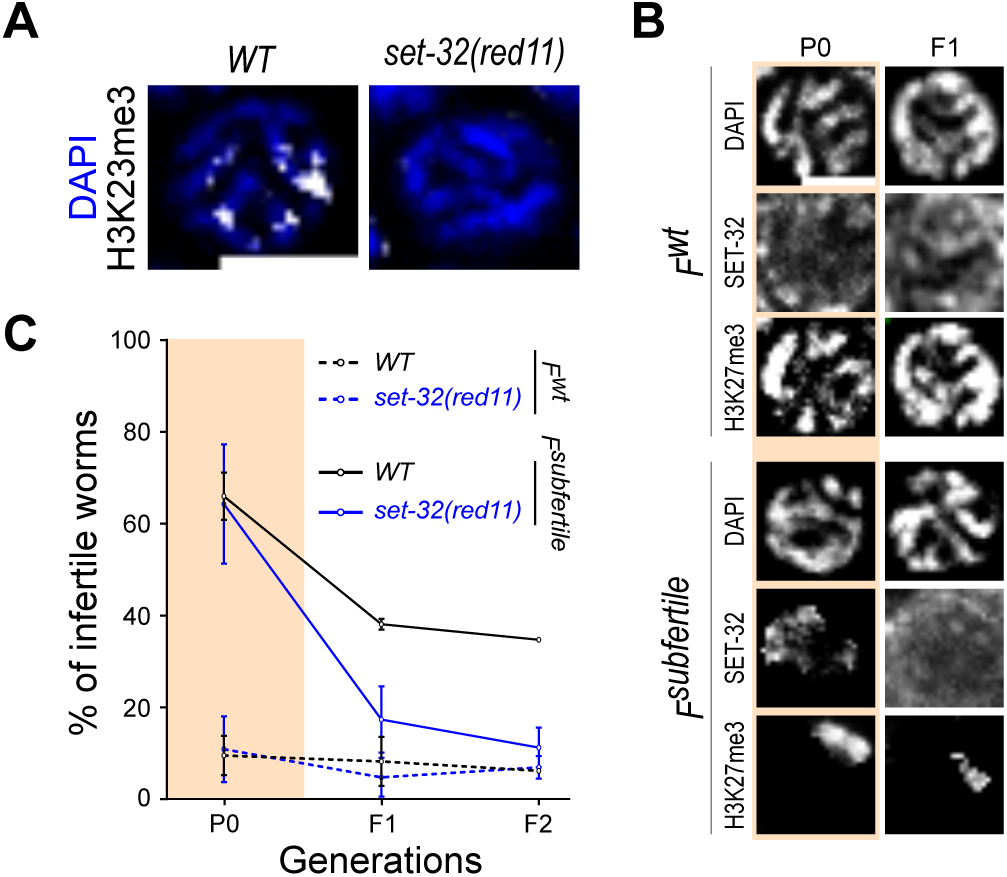
The H3K23 HMT SET-32 is required for initiating the EI of *H3.3K27M^tet^*-induced defects. **(A)** Representative images from immunostaining of H3K23me3 (white) in late pachytene nuclei of wildtype and *set-32(red11)* animals grown at 25°C. **(B)** Representative images from immunostaining of H3K27me3 and FLAG-tagged SET-32 in late pachytene nuclei of *P0-F1^wt^* and *P0-F1^subfertile^* generations. Scale bars represent 4 µm in **(A and B)**. **(C)** Percentages of infertile worms in *P0-F2^wt^* (solid lines) and *P0-F2^subfertile^*(dashed lines) generations in wildtype and *set-32(red11)* background at 25°C. Means and standard deviation of at least three biological replicates are shown.

**Table S3.** List of strains and transgenes used in the study.

| Strain | Genotype | Source | Purpose |
| --- | --- | --- | --- |
| N2 | wildtype | CGC |  |
| FAS158 | *ugeEx108* | This study | Tetracycline inducible multi copy extrachromosomal *H3.3K27M* |
| ALS132 | *cec-6(ele12::loxP+) IV* | Saltzman et al. (2018) | *cec-6* KO |
| FAS213 | *cec-6(ele12::loxP+) IV; ugeEx108* | This study | *cec-6* KO with tetracycline inducible multi copy extrachromosomal *H3.3K27M* |
| FAS233 | *ugeTi144; jsTi1493ugeSi115* | This study | Tetracycline inducible single copy integrated *H3.3K27M* |
| FAS264 | *ugeTi144; cec-6(ele12::loxP+) IV; jsTi1493ugeSi115* | This study | *cec-6* knockout with tetracycline inducible single copy integrated *H3.3K27M* |
| ALS611 | *cec-6(ele23[cec-6::2xTEV::3xFLAG + loxP]) IV* | This study | FLAG tagged endogenous *cec-6* |
| FAS268 | *cec-6(ele23[cec-6::2xTEV::3xFLAG + loxP]) IV; ugeEx108* | This study | FLAG tagged endogenous *cec-6* with tetracycline inducible multi copy extrachromosomal *H3.3K27M* |
| YY1481 | *heri-1::gfp::3xflag II* | Perales et al. (2018) | FLAG tagged endogenous *heri-1* |
| FAS259 | *heri-1::gfp::3xflag II; ugeEx108* | This study | FLAG tagged endogenous *heri-1* with tetracycline inducible multi copy extrachromosomal *H3.3K27M* |
| CSS186 | *set-32(red11)I* | Kalinava et al. (2018) | SET domain deletion of *set-32* |
| FAS236 | *set-32(red11)I; ugeEx108* | This study | SET domain deletion of *set-32* with tetracycline inducible multi copy extrachromosomal *H3.3K27M* |
| FAS284 | *uge157ele23* | This study | Chromodomain dead mutation of *cec-6* |
| FAS285 | *uge157ele23; ugeEx108* | This study | Chromodomain dead mutation of *cec-6* with tetracycline inducible single copy integrated *H3.3K27M* |
| FAS286 | *uge158[heri-1(W62A,Y65A)::GFP::3xFLAG]* | This study | Chromodomain dead mutation of *heri-1* |
| FAS287 | *uge158[heri-1(W62A,Y65A)::GFP::3xFLAG]; ugeEx108* | This study | Chromodomain dead mutation of *heri-1* with tetracycline inducible single copy integrated *H3.3K27M* |

| Transgenes | Genotype | Purpose |
| --- | --- | --- |
| *ugeEx108* | *rpl-28p::rTetR::QF::P2A::mKate::T2A::TetR::pie-1 + 7xTetO::pes-10p::his-72[K27M]::ollas his-72* | Driver and responder transgene |
| *ugeTi144* | *rpl-28p::rTetR::QF::P2A::mKate::T2A::TetR::pie-1+ unc-119 (+) II* | Driver transgene |
| *jsTi1493ugeSi115* | *7xTetO::pes-10p::his-72[K27M]::ollas::his-72 3'UTR IV* | Responder transgene |
| *uge157ele23* | *cec-6(uge157ele23[cec-6(W55A,Y58A)::2xTEV::3xFLAG + loxP]) IV* | Chromodomain dead mutation of *cec-6* |
| *uge158* | *heri-1(W62A,Y65A)::GFP::3xFLAG* | Chromodomain dead mutation of *heri-1* |

**Table S5.**
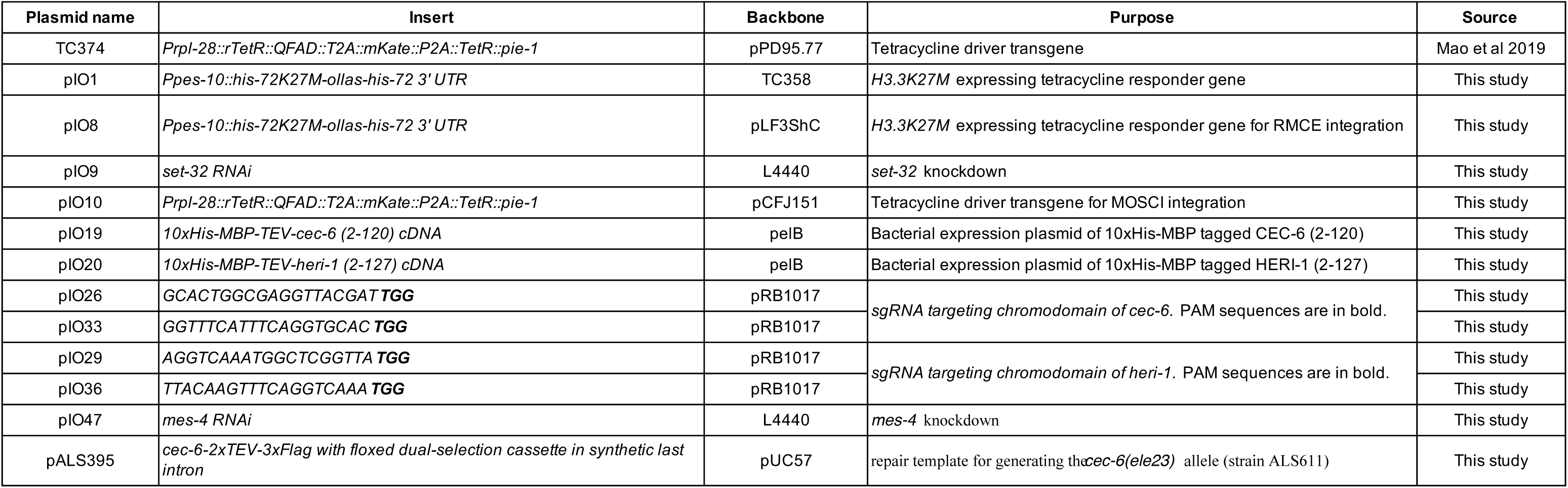
List of plasmids used in the study.

**Table S6.** The fixation conditions for immunofluorescent experiments.

| Primary Ab | Co-staining primary Ab | Fixation procedure |
| --- | --- | --- |
| H3K27me2/3 | OLLAS | 20 minutes methanol +10 minutes acetone |
|  | FLAG | 5 minutes ethanol +5 minutes acetone |
|  | H3K36me3 | 10 minutes methanol +5 minutes acetone |
|  | H3K23me3 | 5 minutes methanol +5 minutes acetone |

**Table S7.** The list of histone H3 peptides used in the study.

| The name of peptide | peptide sequence |
| --- | --- |
| H3 | *Bio*-KAPRKQLATKAARKSAPASGGK |
| H3K27M | *Bio*-KAPRKQLATKAARMSAPASGGK |
| H3K9me3 | *Bio*-ARTKQTARK(me3)STGGKAPRKQL |
| H3K23me3 | *Bio*-KAPRKQLATK(me3)AARKSAPASGGK |
| H3K27me3 | *Bio*-KAPRKQLATKAARK(me3)SAPASGGK |
| H3K36me3 | *Bio*-AARKSAPASGGVK(me3)KPHRYRP |

*Bio*- N-terminal biotin tag
K27M mutation and trimethylation are highlighted in red

